# How brain pulsations drive solute transport in the cranial subarachnoid space: insights from a toy model

**DOI:** 10.64898/2026.02.24.707684

**Authors:** Alannah Neff, Alexandra Vallet, Mariia Dvoriashyna

**Affiliations:** School of Mathematics and Maxwell Institute of Mathematical Sciences, University of Edinburgh, Edinburgh, UK; Mines Saint-Etienne, INSERM, U1059 SAINBIOSE, F42023, Saint-Etienne, France

**Keywords:** Cerebrospinal fluid, glymphatic system, transport in oscillatory flows, steady streaming, lubrication theory

## Abstract

Cerebrospinal fluid (CSF) circulates around and through the brain, supporting neural homeostasis by regulating the extracellular chemical environment. Yet the physical mechanisms governing CSF-driven solute transport remain poorly understood, limiting the design of diagnostic and therapeutic strategies targeting brain clearance and drug delivery.

Pulsatile CSF flow in the cranial subarachnoid space (cSAS), is driven by cardiac, respiratory, and sleep-related vasomotion. Over longer timescales weaker steady flows, such as inertial steady streaming, Stokes drift, and production–drainage flow, may contribute to solute transport, but their role and relative importance remain unclear.

Here, we develop a simplified two-dimensional model of CSF flow and solute transport in the cSAS using lubrication theory. Through multiple-timescale and asymptotic analyses, we derive a reduced long-time transport equation in which advection is governed by the Lagrangian mean velocity, incorporating steady streaming, production–drainage flow, and Stokes drift.

Analysing three physiologically relevant case studies, we show that steady flows can substantially reshape concentration profiles, enhance dispersion, and alter clearance efficiency. Our results clarify the mechanisms underlying CSF-mediated transport, predict distinct regimes in humans and mice, and highlight the importance of subject-specific physiological parameters when interpreting contrast-agent and intrathecal drug-delivery studies.

## 1 Introduction

Cerebrospinal fluid (CSF) is a Newtonian fluid produced predominantly by the choroid plexus within the brain’s ventricles [24]. From the ventricles, CSF circulates in the subarachnoid space (SAS) around the brain and spinal canal (see figure 1) [24], where it is thought to act as a clearance pathway for metabolic waste in the interstitial brain tissue (parenchyma) [21, 18, 56]. To collect metabolic waste, CSF flows into the brain along the perivascular spaces (PVS), narrow regions surrounding the penetrating veins and arteries. From the SAS, CSF is proposed to clear at various locations [36], including at intracranial drainage sites such as the arachnoid granulations (AG) and through the cribriform plate (shown in figure 1).

**Figure 1.**
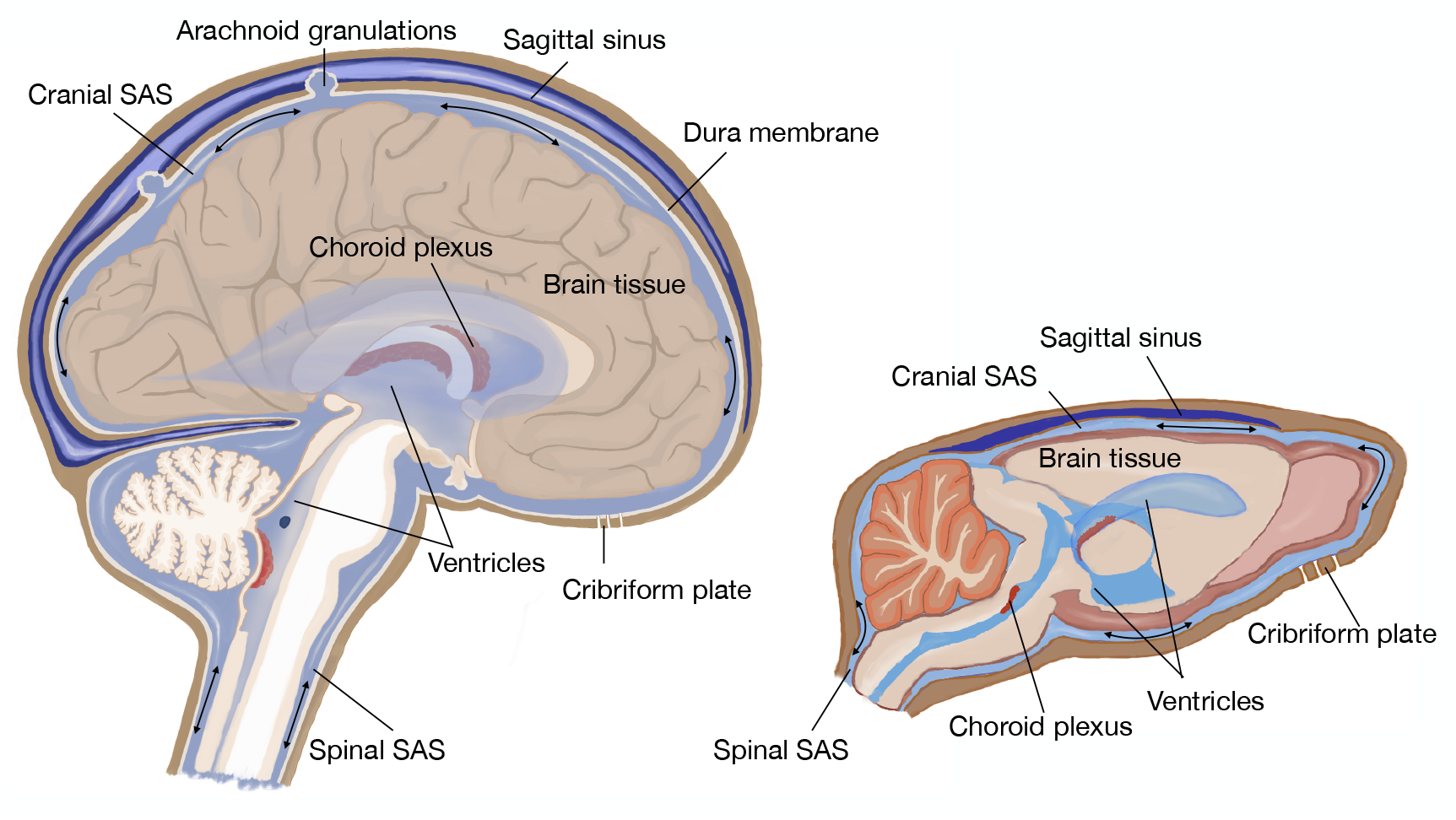
Schematic of the cross-section of the human (left) and mouse (right) brain and cerebrospinal fluid (CSF) compartments. SAS: subarachnoid space. The arrows depict the pulsating flow of CSF in SAS. Drawings were made by MD, drawing of the human brain is based on the image in commons.wikimedia.org.

Disruptions to CSF flow and consequently to solute clearance can lead to the accumulation of proteins such as amyloid-*β* and *α*-synuclein, which are implicated in the pathology of Alzheimer’s disease and Parkinson’s disease, respectively [7, 31]. CSF also provides a route for intrathecal drug delivery, where the drug is injected directly into the CSF in the spinal SAS, therefore circumventing the blood-brain barrier [59]. A deeper understanding of CSF transport pathways is vital to identify the mechanisms contributing to neurological disorders and to support the development of CSF-based therapeutic interventions.

The precise mechanisms by which CSF circulates and clears solutes remain debated. Recent human imaging studies reveal that brain pulsations due to changes in cerebral blood volume throughout the cardiac [1, 46] and respiratory cycles [17, 46, 60] generate oscillatory CSF flow in the SAS and ventricles. Furthermore, slow waves (0.05–0.1 Hz in humans) driven by vasomotion and the contraction and dilation of smooth muscle cells in cerebral blood vessels, increase this oscillatory CSF motion during NREM sleep [20, 52]. Oscillatory displacement of CSF has also been observed in the pial (surface) PVS of mice, induced by arterial wall motion [32] and vasomotion during sleep [19].

Measuring solute transport in the human SAS is challenging. There are measurements of oscillatory CSF dynamics at good temporal resolution, e.g. capturing the dynamics over a cycle of oscillations; however, they are typically confined to a plane such as the aqueduct, foramen magnum or C2/C3 in the spinal canal rather than the full cranial SAS (cSAS) [47, 17, 60]. On the other hand, tracer studies which capture the evolution of a contrast agent throughout the entire central nervous system often have good spatial resolution but are restricted to discrete time points, on a timescale much longer than the period of physiological oscillations [39, 41]. Mathematical modelling has emerged as a relevant tool, which can utilise the available CSF velocity measurements and physical laws to synthesise the fast oscillatory dynamics of CSF with the long-term transport of solutes [24, 11]. Pulsatile CSF flows have been modelled in the spinal SAS, using asymptotic [44, 45, 10] and numerical methods [25, 37, 57], and have also been studied in both a single [51] and a network of PVS [14, 15]. Fewer models exist in cSAS, e.g. a numerical study of CSF flow [16] and CSF flow with drug delivery [49].

Transport of solutes in oscillatory flow occurs as a combination of advection - the fluid carrying the solute - and diffusion. Understanding the interplay between these transport mechanisms can aid interpretation of exploratory imaging studies and the design of drug intervention protocols. In the spinal SAS, analytical modelling showed that oscillatory flows appear in the long-time transport dynamics via Lagrangian mean velocity [44, 27], comprised of steady streaming and Stokes drift. Steady streaming arises from inertial effects due to oscillations [38, 26], and has been shown to play a role in drug delivery for the inner ear [48] and as a mixing mechanism in the anterior chamber of the eye [12]. Stokes drift arises when spatial variations in oscillatory velocity cause fluid particle displacements to accumulate into a net drift over successive cycles [26]. The effects of steady streaming and Stokes drift have remained thus far unexplored for the cSAS.

In this work, we present a simplified 2-dimensional (2D) model of CSF flow and solute transport in the cSAS, with the aim of identifying relevant transport mechanisms. The purpose of the model is to understand the relative importance, depending on the physiological phenomena generating the oscillatory flow, of physical mechanisms for the transport of solutes in CSF. In conjunction with oscillatory flows, a small net CSF flow may exist due to the production and drainage of CSF. One proposed route stems from the production of CSF in the choroid plexus in the ventricles and drainage of CSF via intracranial absorption routes [36], which we incorporate into our model. Our model provides a first step towards the development of an analytical model of solute transport around the brain, complementing existing numerical works and providing an efficient way to explore the role of physiological parameters on solute transport.

This paper is structured as follows. In section 2, we first present a simplified 2D model of CSF flow and solute transport in the cSAS. We use lubrication theory, an asymptotic technique that takes advantage of the small aspect ratio of the domain, *ε*. Following a scaling analysis, in section 2a, we identify that for human oscillations, we expect secondary steady flows to play a leading role in solute transport. We solve analytically at successive orders of *ε* for the leading-order oscillatory and higher-order steady flows. Following the procedure in [27] we derive a mean transport equation, averaged over the timescale of oscillations in section 2b. We then solve this equation for three physiological case studies in section 3. We use our framework to assess the relative importance of steady streaming, Stokes drift and production–drainage flows for the transport of solutes in the cSAS. Finally, we discuss the implications for transport in mice and humans in section 4.

## 2 Methods

We consider a model of fluid flow and resulting solute transport in the cSAS. CSF is modelled as an incompressible, Newtonian fluid with motion governed by the Navier–Stokes equations. The geometry of the cSAS is simplified to a long, thin channel, of length *L* (the circumference of the brain) and time-dependent thickness 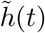, such that the domain spans 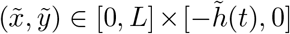, under the assumption that the inverse curvature of the domain is much larger than its thickness. CSF flow is primarily driven by periodic pulsations of the brain, such as cardiac, respiratory, or slow-wave oscillations, of distinct amplitude *Ã* and frequency *f*. These are modelled as a transverse, time-periodic oscillation of the lower boundary of the channel, described by 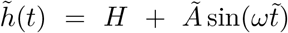, where *ω* = 2*πf* denotes the angular frequency of the oscillations. We model the boundary motion as a single sinusoidal oscillation to isolate the contribution of each forcing type to solute transport. In reality, the boundary motion occurs due to a combination of physiological oscillations, which may be incorporated into the model as a superposition of sinusoidal modes at different -frequencies.

The transport of solutes such as drugs and metabolites in CSF occurs as a combination of advection by fluid motion and molecular diffusion. In this 2-D geometry, a solute concentration 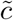 in the cSAS evolves according to the advection–diffusion equation

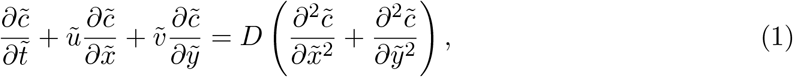

where *D* is the diffusion coefficient of the solute, and (*ũ*, 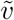) is the velocity of CSF. Representative dimensional parameter values, summarised in table 1, are obtained from the physiological parameter estimation detailed in Supplementary Material section S10, where relevant sources are cited.

**Table 1:**
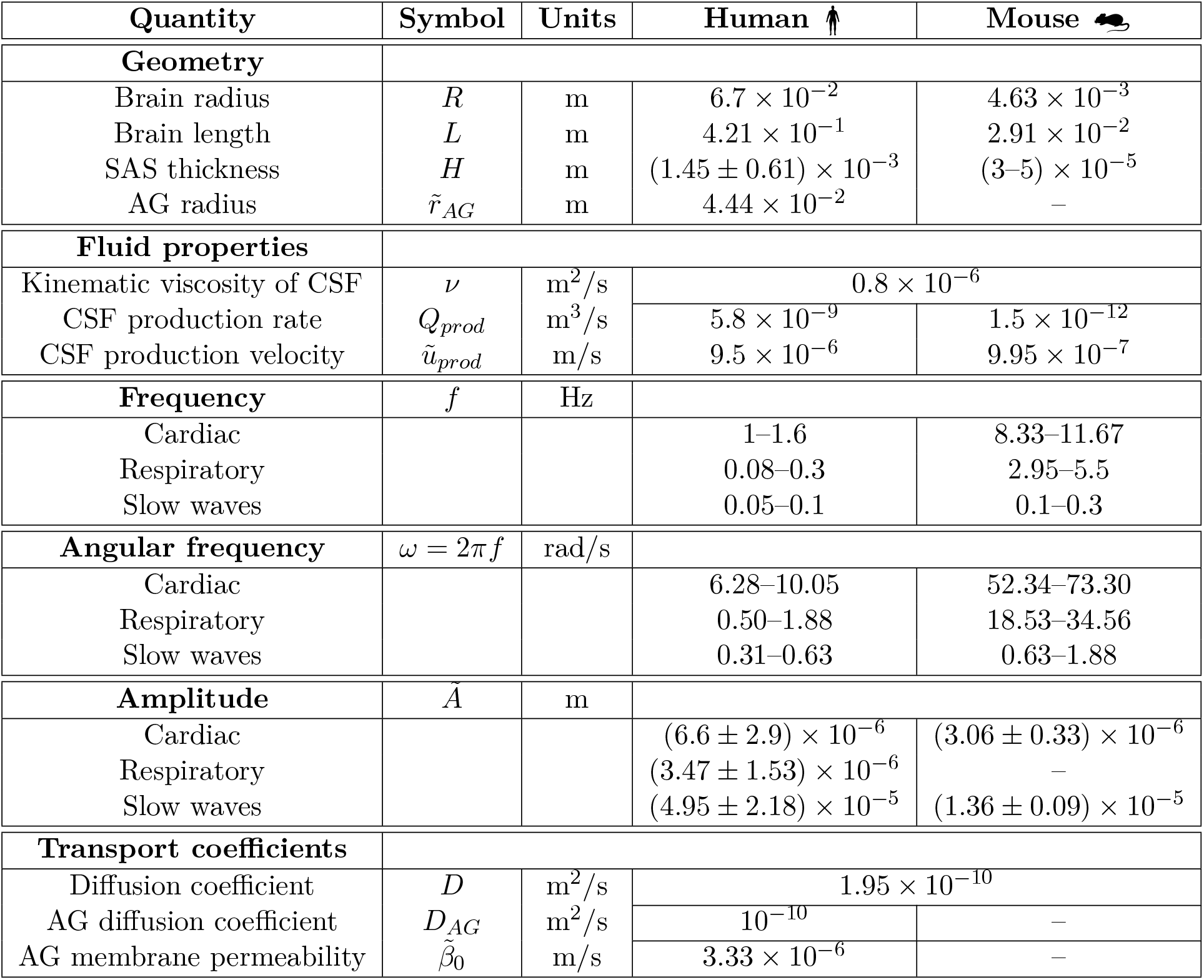
Physiological parameters and fluid properties. Note: ± denotes a mean and standard deviation, while − corresponds to a range of values. The justification for all values and corresponding references are reported in Supplementary Material section S10.

### 2a Timescale Analysis

Motivated by classifying the solute transport regime induced by brain pulsations, we perform a timescale analysis of the advection–diffusion equation equation (1). For the channel domain in Cartesian coordinates and fluid velocity ***ũ*** = (*ũ*, 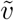) we posit lubrication-type scalings such that

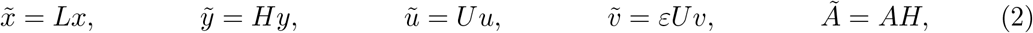

where the dimensional variables are denoted with tildes. The small aspect ratio gives rise to the dimensionless parameter *ε* = *H/L* which is calculated to be *ε* = 0.003 from the values in table 1. The dimensionless amplitude *A* is *O*(*ε*), since the mean amplitudes in table 1 are an order of magnitude smaller than the corresponding SAS thickness, such that *A*_0_ = *A/ε* ∼ *O*(1). We scale the vertical velocity by the amplitude and frequency of oscillations, *Ãω*, and the horizontal velocity scales as *U* = *Ãω/ε* (details in Supplementary Material section S2).

The natural timescale of the problem, *t*_N_, is *ω*^−1^. To characterise the possible transport regimes arising in oscillatory CSF flows, we consider the three additional timescales within this problem. First, the timescale of transversal diffusion is 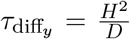, where *D* is the molecular diffusion coefficient of the solute. The second timescale is that of advection as a result of oscillatory flow 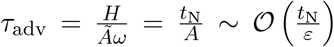. The final timescale corresponds to longitudinal diffusion 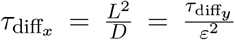, which is two orders of magnitude slower than diffusion in the transverse direction.

To classify the different transport regimes, we consider the Péclet number

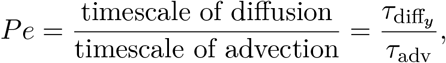

of the leading advective and diffusive mechanisms. Using the definition above, 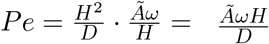.

Depending on the value of *Pe*, we identified the following possible transport regimes:

1. *Pe* ≪ *ε*^2^**: Diffusion dominated**. In a diffusion-dominated regime, longitudinal diffusion, the slower of the two diffusive mechanisms, occurs much faster than the timescale of advection, such that 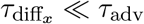.
2. *Pe* ∼ *ε*^2^**: Mixed transport**. Due to advection and longitudinal diffusion occurring on the same timescale 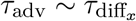, transport is attributed to a combination of the two. The concentration profile in *y* is spatially homogeneous since the transverse diffusion occurs on a shorter timescale than the characteristic timescale of the problem *t*_N_.
3. *Pe* ∼ *ε***: Taylor dispersion**. When transverse diffusion occurs on the timescale of the oscillations driving the flow, *t*_N_, such that 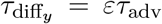, we get the Taylor-dispersion regime [45]. The concentration profile is homogeneous across the *y*-direction, and the transport can be described by effective diffusion and by advection [50, 22].
4. *Pe* ∼ 1**: Mixed transport**. Due to transverse diffusion and advection occurring on the same timescale, 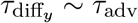, solutes are transported through their combination.
5. *Pe* ≳ 1*/ε***: Transport by secondary flows and transverse diffusion**. In this case *τ*_adv_ *< τ*_diff_, and solute transport occurs via oscillatory flow. If secondary flows such as steady streaming and Stokes drift exist, these may occur on the same timescale as transversal diffusion, contributing to long-time transport.

To examine the effect of parameter variation on the resulting transport regime, we write the Péclet number, *Pe* = *Aα*^2^*S*, in terms of dimensionless amplitude *A* = *Ã/H*, the Womersley number *α*, defined as *α*^2^ = *ωH*^2^*/ν* where *ν* is the kinematic CSF viscosity, taken as constant (table 1), and the Schmidt number *S* = *ν/D. α* and *S* quantify the relationship between kinematic viscosity and oscillatory inertia, and kinematic viscosity and molecular diffusivity, respectively. Dimensionless groups and their values are reported in table 2. *Pe* characterises the transport regime based on the interactions between the oscillatory dynamics, fluid properties, and solute diffusivity. If multiple oscillatory components were considered with distinct amplitudes and frequencies, their combined effect would modify the advective transport and hence the corresponding Péclet number.

**Table 2:**
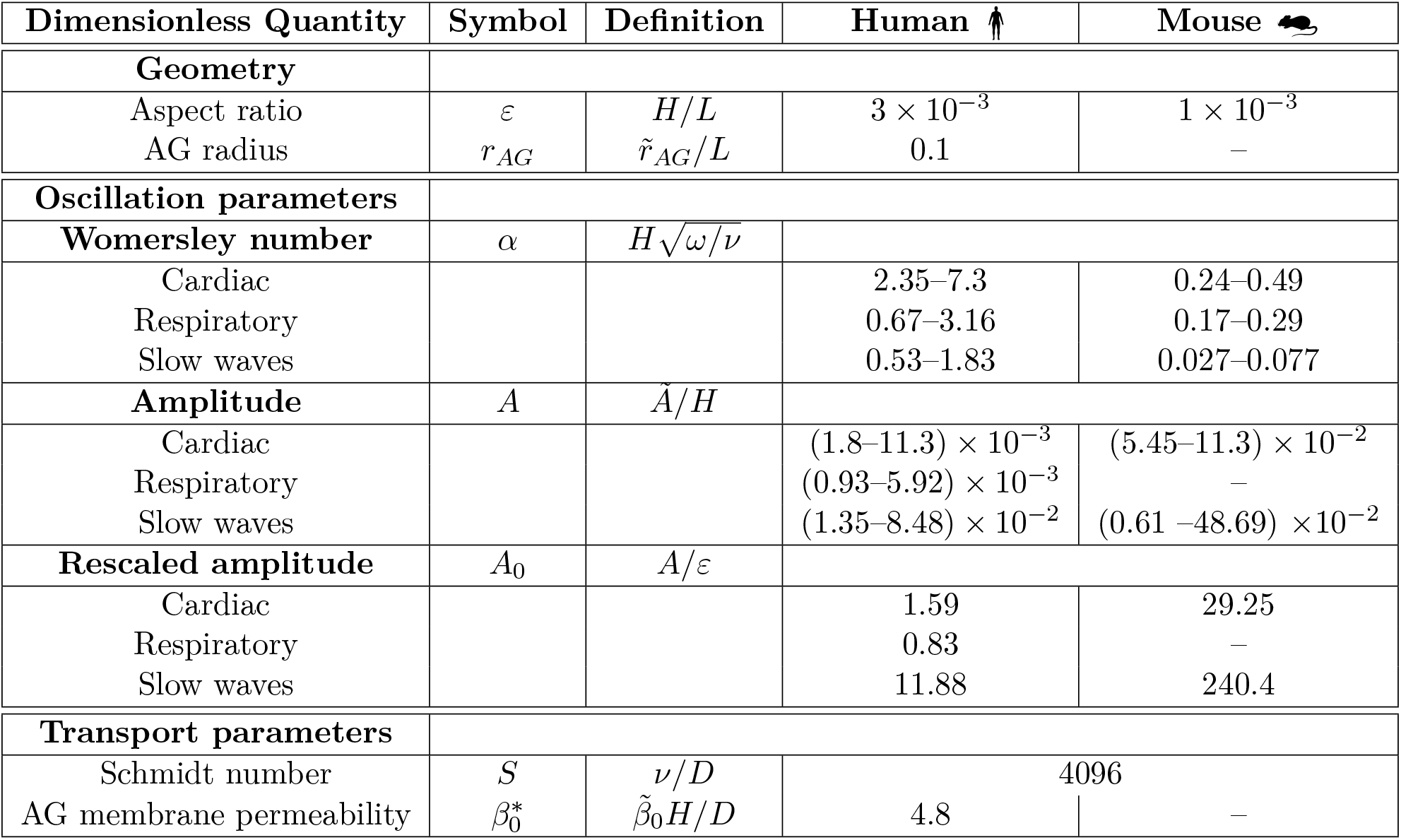
Dimensionless parameters used in the model. Parameters are calculated for the values in table 1 with details provided in Supplementary Material section S10. *A*_0_ are calculated for mean *A*.

The molecular diffusivities of relevant solutes lie within the same order of magnitude, e.g. amyloid-*β* (*D* ≈ 1.8 × 10^−10^ m^2^/s [58]) and gadobutrol (*D* ≈ 3.8 × 10^−10^ m^2^/s [53]). We therefore select a representative diffusivity *D* (see table 1) within this range, giving a Schmidt number *S* = 4096 ∼ *O*(*ε*^−2^).

With *S* fixed, we plot the Péclet number in figure 2 across a range of physiological values of *A* and *α* arising from species differences, individual variability, and physiological states. For the parameter values associated with cardiac and vasomotion oscillations in the human brain listed in table 1, solutes in CSF will be transported in regimes 4—5. Below, we focus on the transport dynamics in regime 5, however the following derivation of a long-time transport equation for solutes transported by CSF also captures the transport dynamics in regime 4 [27].

**Figure 2.**
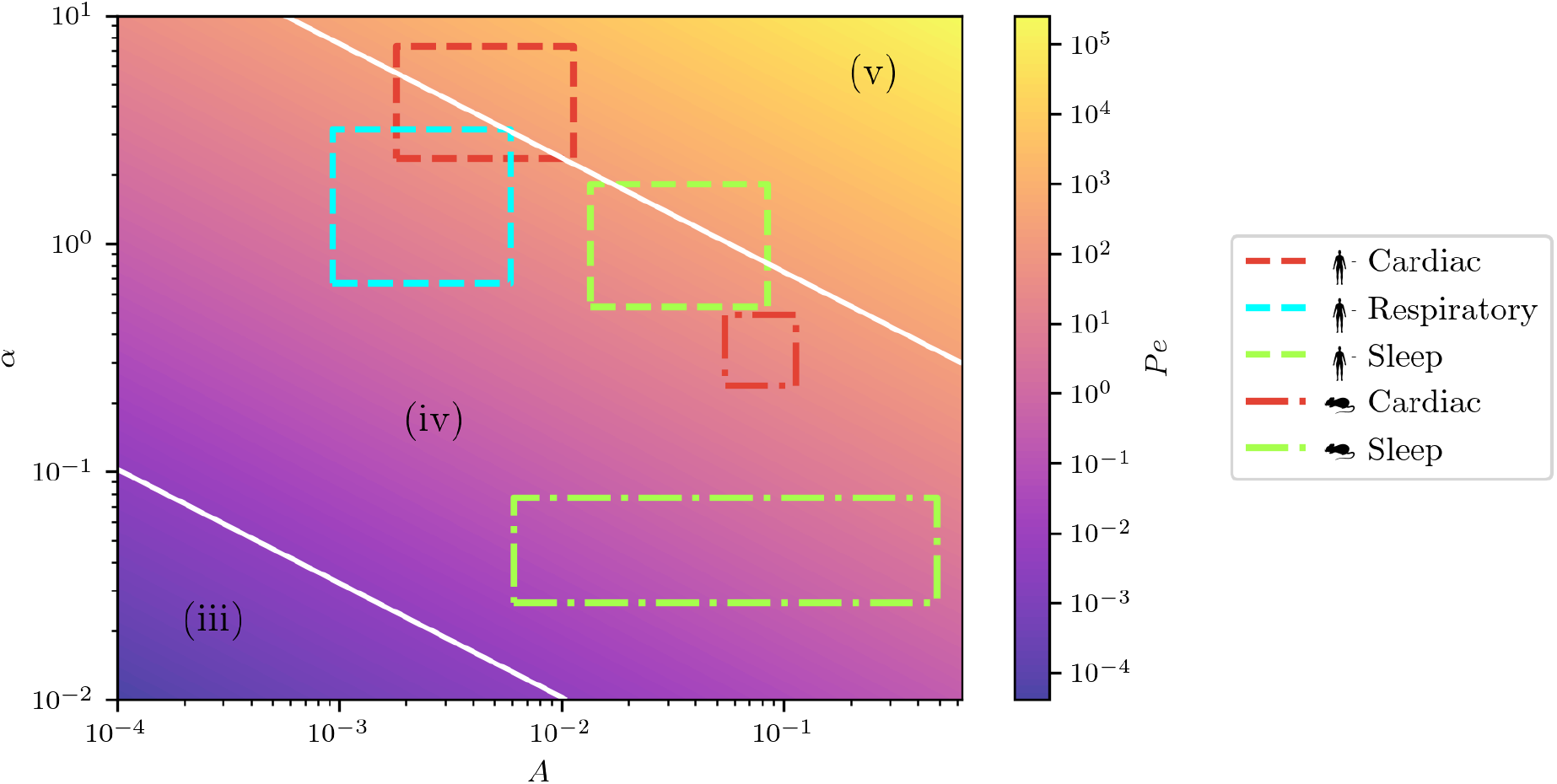
Transport regimes associated with oscillatory flow in a channel with a uniformly oscillating lower boundary. The transport regimes associated with pulsations driven by the cardiac cycle (red), respiration (cyan) and sleep (green) are outlined for humans (dashed) and mice (dot-dashed). White lines indicate approximate boundaries between regimes 3–5. Transport by brain pulsations occurs mostly around transport regime 4–5, where advection by secondary flows is important. Despite these oscillations having different amplitudes and frequencies, each lies within the regime where the rate of advection is approximately an order of magnitude larger than the rate of transversal diffusion. Pulsations generated by respiration in humans, and both cardiac and vasomotion pulsations in mice are in regime 4, transported by mixed transport. In each of these regimes, diffusion in the longitudinal direction is negligible compared to other transport mechanisms. We fix *S* = 4096 from the values of *D* and *ν* in table 1. To obtain a similar figure for a different solute, one must recalculate *Pe* with the corresponding *S*. For larger solutes, such as gadobutrol and amyloid-*β* we expect similar behaviour. However, for small ions, the slow-wave and respiratory pulsations fall between regimes 4 and 5 for humans. Regimes 1 and 2 (not shown) are not reached by physiological oscillations.

### 2b Model of fluid and solute transport

As advection plays an important role in regimes 4–5, the CSF velocity field must be found before using it in the advection–diffusion equation to study solute transport. The non-dimensionalised Navier–Stokes equations for an incompressible fluid and the solute transport equation are

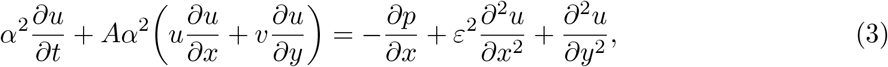

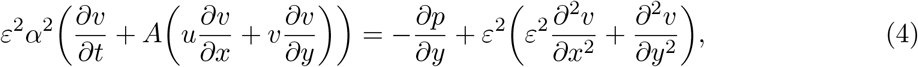

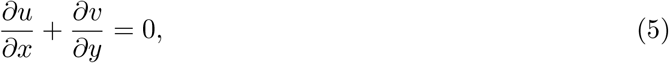

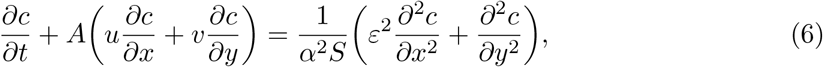

where ***u*** = (*u, v*) is the fluid velocity, *p* is the fluid pressure and *c* is the concentration of solute. The non-dimensionalisation of the equations is detailed in Supplementary Material section S2. Equations (3) and (4) describe conservation of momentum for the fluid, equation (5) conserves the fluid mass and equation (6) is the advection–diffusion equation describing conservation of solute mass. We note that longitudinal diffusion is of order *ε*^2^, and we will neglect it in this work together with other terms of order *ε*^2^.

We consider flows driven by oscillations of the lower wall and due to the production and drainage of CSF. We model the latter by introducing a constant outflow in a generic absorption site located at the centre of the dura membrane. We focus on production–drainage flow in humans and assume drainage through arachnoid granulations, structures which protrude through the dura mater, connecting the cSAS with the venous sinuses. CSF production by the choroid plexus occurs at the rate *Q*_*prod*_ = 5.8 × 10^−9^ m^3^/s. The expected velocity of production–drainage flow *ũ*_*prod*_ = *Q*_*prod*_ */*(2*πRH*) ^≈^ 9.5· 10^−6^ m/s, is about *ε* times smaller than that of the oscillatory flow *U* = *Ãω/ε* ^≈^ 1.2 ·10^−2^ m/s during the cardiac cycle (in humans). We therefore model CSF drainage at the upper boundary *y* = 0, by prescribing a dimensionless normal velocity of the form *v* = *εq*(*x*) = *εq*_0_*d*(*x*), where

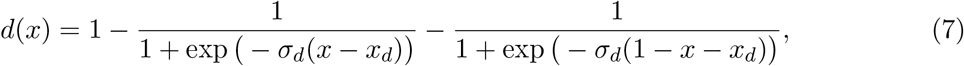

which represents a localised outflow centred at *x*_*d*_ with steepness controlled by *σ*_*d*_. The magnitude *q*_0_ is determined by the rate of CSF production through

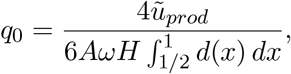

so that the resulting flow corresponds to a balance between production and drainage. The boundary conditions for the fluid are summarised in figure 3. Each case study in section 3b will have specific solute boundary conditions.

**Figure 3.**
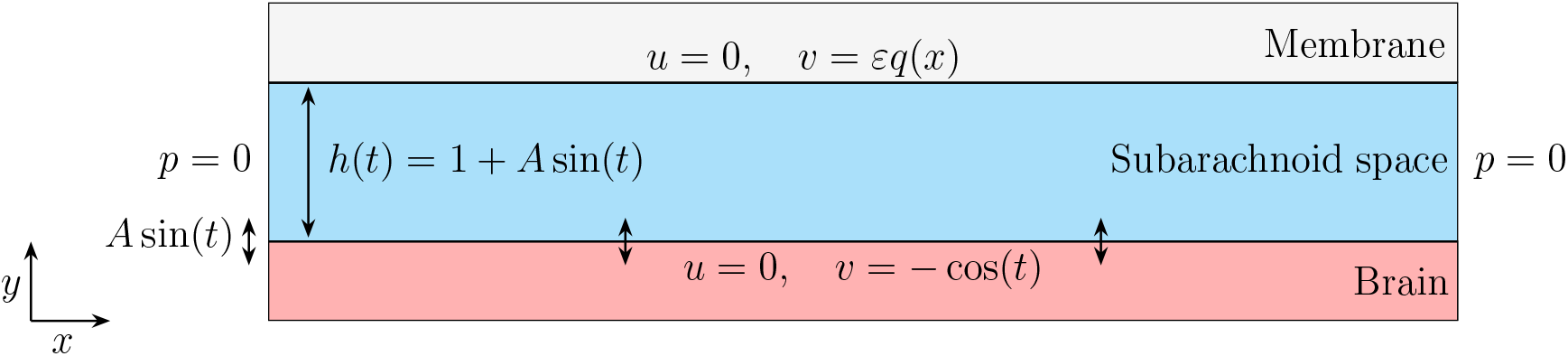
Summary of the (dimensionless) fluid flow problem associated with the flow of CSF in the cSAS driven by physiological pulsations of the brain. The brain is considered an impermeable membrane with a normal velocity equal to the velocity of the oscillating boundary. At both the brain and the dura membrane, we assume a no-slip boundary condition. We assume that the fluid can exchange freely between the cranial and spinal SAS from the ends of the channel (*x* = 0, 1), and prescribe *p* = 0 there. At the dura membrane (at *y* = 0) we prescribe the outflow velocity ***u*** = (0, *εq*(*x*)). The justification for *q*(*x*) and the corresponding parameters are given in section S10(c).3. The magnitude of *q*(*x*) is dependent on the rate of CSF production *Q*_*prod*_.

It is convenient to work in the fixed domain, thus we perform a change of variables *η* = *y/h*(*t*). This change of variables introduces an additional transverse inertial term to equations (3) to (6). In Supplementary Material section S4 we report the detailed asymptotic expansion in powers of *ε* of the above system.

We expand the velocity and pressure components asymptotically and separate each term into its harmonics. Firstly, we solve the linear leading-order problem for the oscillatory flow ***u***_0_, detailed in section S4(a). At the following order, the steady, time-independent flow ⟨***u***_1_⟩ is derived by solving the short-time-averaged 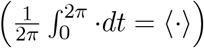, nonlinear, *O*(*ε*) problem. The steady flow is comprised of the steady streaming and production–drainage flows such that ⟨***u***_1_ ⟩ = ***u***_*s*_ + ***u***_*pd*_ (see equations (S36), (S37), (S43) and (S44)).

On the timescale of oscillations, the solute is simply advected by the leading-order oscillatory flow, resulting in no net transport of the solute as its profile oscillates with the flow (section S5). However the secondary flows occur on the same timescale as transversal diffusion. To quantify the advective contribution of secondary flows, we derive an expression for the long-time fluid particle trajectories, also known as the Lagrangian mean velocity. The fluid particle trajectories are influenced by both the steady flows ⟨***u***_1_⟩ and contributions from the oscillatory flow known as Stokes drift ***u***_*ω*_. Particle trajectories in the (*x, η*)-domain are defined by

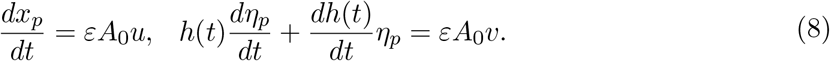

The timescale of the long-time fluid particle trajectories *τ* ∼ (*ε*^2^*A*_0_*ω*)^−1^ = (*ε*^2^*A*_0_)^−1^*t*, where relative displacements of *O*(1) occur, is obtained by performing a dimensional analysis on equation (8). We exploit the two timescales in the problem to find the leading-order particle trajectories by a multiple-timescale expansion.

The variables *x*_*p*_ and *η*_*p*_ are expanded asymptotically in *ε* where each component is assumed to be periodic in *t*. We show in Supplementary Material section S5 that the leading-order particle positions satisfy the following long-time dynamics equations

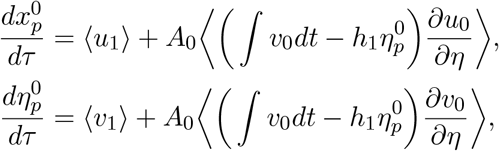

where ***u***_0_ and ***u***_1_ denote leading and (*ε*) velocity solutions, and *h*_1_(*t*) = *A*_0_ sin *t* arises as the *O*(*ε*) term from the asymptotic expansion of the sinusoidal oscillation *h*(*t*). The Lagrangian mean velocity is defined as

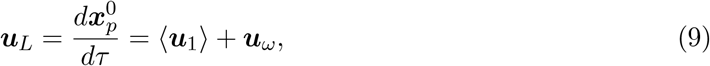

where ⟨***u***_1_⟩ = ***u***_*s*_ + ***u***_*pd*_ is the summation of the steady streaming and production–drainage flow, and ***u***_*ω*_ is the Stokes drift of the fluid particle.

The long-time transport equation is derived by performing a multiple-timescale expansion of the advection–diffusion equation in *t* and *τ* (details in section S7). Once more leveraging asymptotic expansions, and solving the *O* (1), *O*(*ε*) and *O* (*ε*^2^) problems sequentially, we obtain the long-time transport equation for leading-order concentration *c*_0_

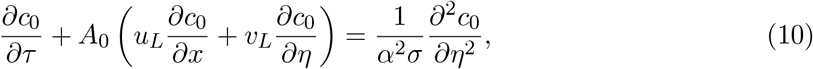

where *σ* = *ε*^2^*S*. The long-time transport equation describes the net transport of a solute in CSF, averaged over the period of oscillations.

### 2c Numerical method

We solve the time-dependent equation (10) for the scalar solute concentration field *c*_0_(*x, η, t*) in a rectangular domain using the finite element method (FEM) implemented in FEniCS [3, 29]. The velocity field ***u***_*L*_ is obtained from the analytical solutions to the fluid flow derived in sections 2b, S4(a), S4(c) and S4(d).

The domain is discretised using second-order Lagrange finite elements. The velocity field is projected onto the same finite-element space. The diffusion coefficient in the *η* − direction is set to be *ε*^4^ smaller than *D*_*x*_ such that the effect of longitudinal diffusion is removed as in the time-averaged transport equation (10). We retain this small longitudinal diffusion coefficient for numerical stability. We employ an implicit backward Euler scheme for time integration. The mesh resolution (*n*_*x*_, *n*_*y*_) = (100, 200) and time step Δ*t* = 0.001 were chosen based on the convergence analysis reported in Supplementary Material section S8.

The variational form is assembled using FEniCS, and the resulting linear systems are solved with a direct solver (MUMPS). The framework supports general boundary condition configurations. The steady-state solutions are computed using the same FEM framework.

## 3 Results

In this section, we present results for fluid flow and solute transport. The following results are for human physiology, and oscillatory flow generated by cardiac oscillations. We therefore set *ε* = 0.003, where *L* is from table 1, *H* is the mean value from the range in table 1. We fix *α* = 4.5 for convenience, corresponding to a heartbeat frequency of 1.2 Hz, unless otherwise specified. Additionally, we set *S* = 4096.

### 3a Fluid flow

In the following, we present and analyse the velocity profiles of the secondary flows: steady streaming ***u***_*s*_, Stokes drift ***u***_*ω*_, and production–drainage ***u***_*pd*_, which comprise the Lagrangian mean velocity equation (9). Figure 4a shows the steady-streaming flow profile, ***u***_*s*_. CSF flows into the SAS from the spinal canal in the middle region of the channel (around *η* = − 1*/*2), and exits back into the spinal SAS close to the brain surface and dura membrane. The net flow across the channel is zero.

**Figure 4.**
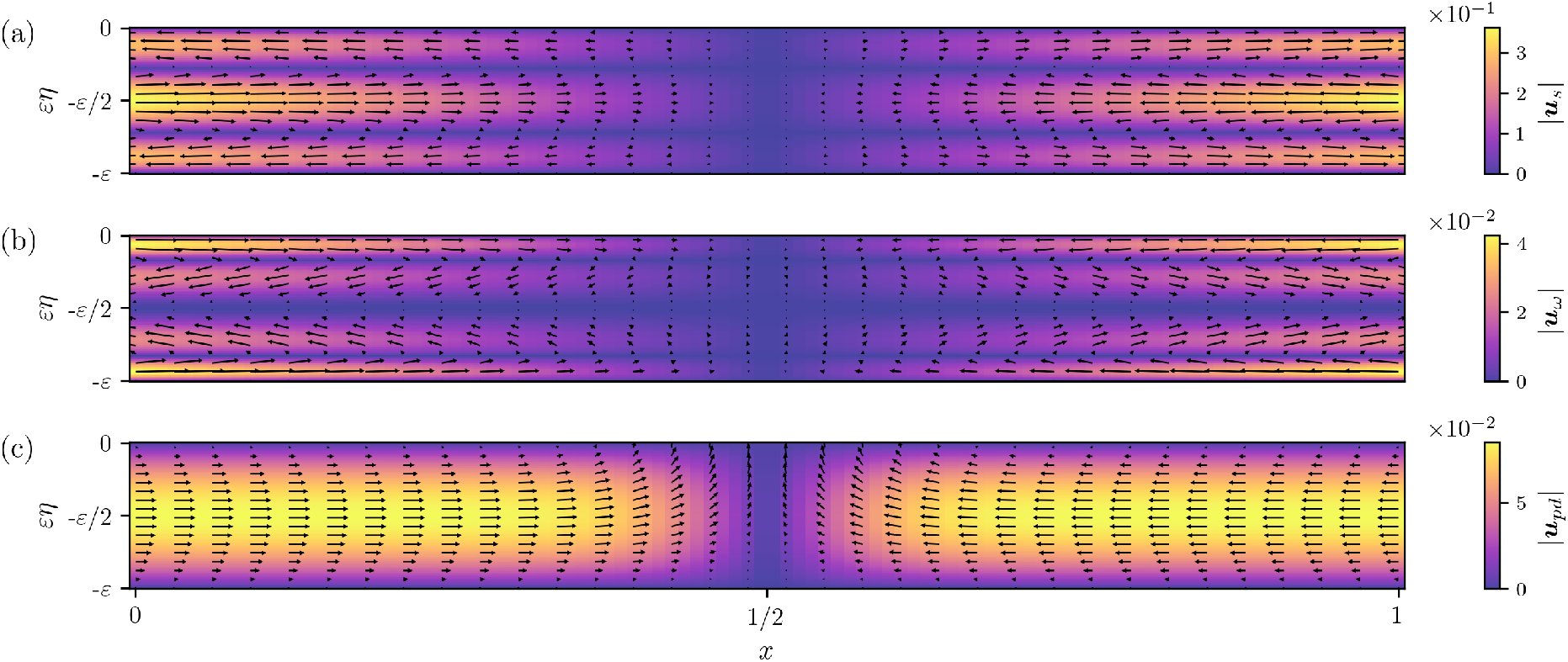
Velocity flow profiles for (a) steady streaming ***u***_*s*_, (b) Stokes drift ***u***_*ω*_, and (c) production–drainage flow ***u***_*pd*_. The colours represent the velocity magnitude, and the arrows are velocity vectors. The thickness of the domain and the vertical velocity components are magnified by a factor of 10 for better visualisation. *A* = 0.01, *α* = 4.5, *ε* = 0.003.

The Stokes drift ***u***_*ω*_ emerging from the leading-order oscillatory flow is shown in figure 4b. CSF flows into the channel in narrow regions close to the brain surface and dura membrane, and exits along the centre of the channel. Production–drainage flow ***u***_*pd*_, shown in figure 4c, exhibits a parabolic-like profile with an outflow region near *x* = 1*/*2 on the upper boundary (*η* = 0), corresponding to drainage through intracranial routes. For the parameter regime considered in figure 4, the steady streaming has the highest magnitude. Moreover, |***u***_*s*_| is an order of magnitude larger than |***u***_*ω*_|. The associated Lagrangian mean velocity ***u***_*L*_ will therefore be dominated by ***u***_*s*_ and ***u***_*pd*_.

Figure 5a shows the (dimensional) maximum values of horizontal components of the ***u***_*s*_, ***u***_*ω*_, ***u***_*pd*_ and ***u***_*L*_ for varying dimensionless amplitude *A*. Stokes drift and steady streaming both vary as *A*^2^, therefore Stokes drift remains an order of magnitude smaller than steady streaming for all reasonable *A*, as shown in figure 4. A similar analysis in *α* in figure S4 shows that both flows are proportional to *α*^4^ for *α <* 10. This suggests that in our problem, invariably, the role of Stokes drift in the transport of solutes is negligible.

**Figure 5.**
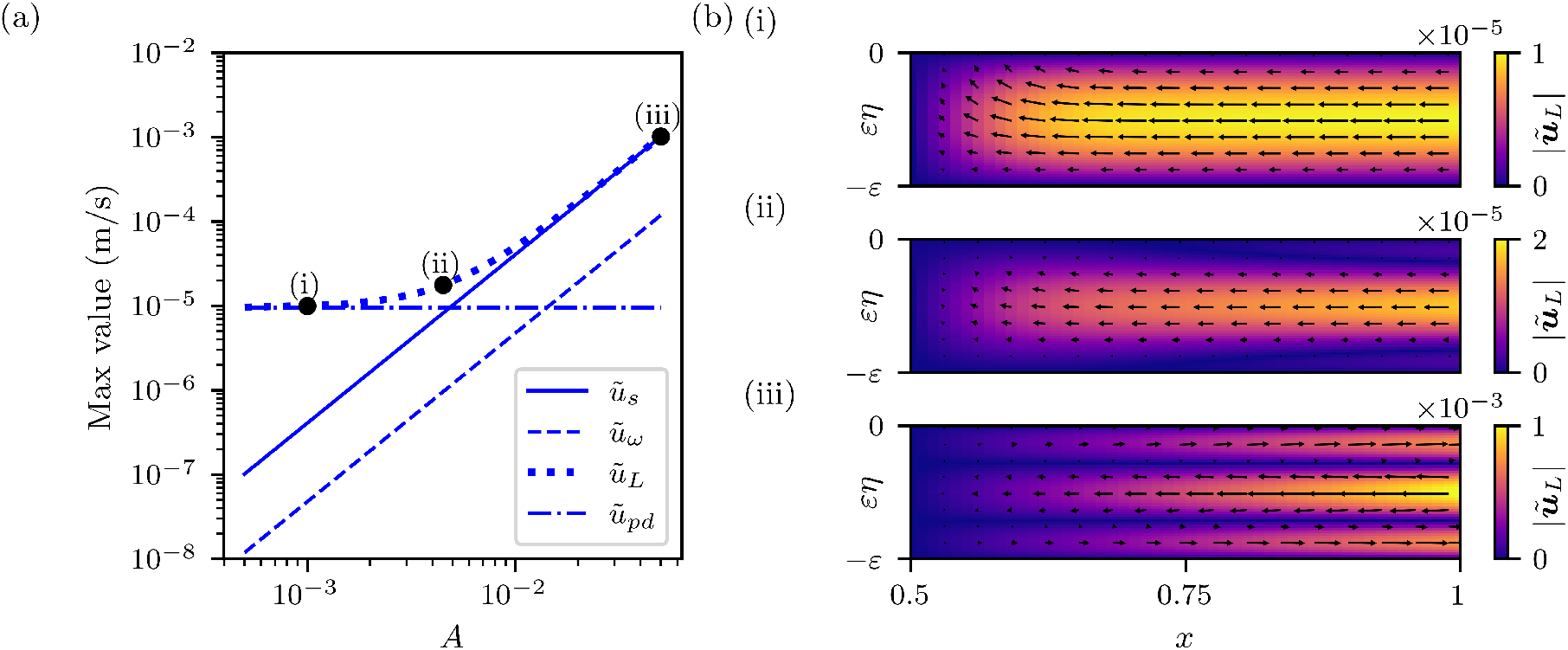
(a) Maximum magnitudes of the dimensional horizontal component of the steady streaming (*ũ*_*s*_), Stokes drift (*ũ*_*ω*_), production–drainage (*ũ*_*pd*_), and Lagrangian mean velocity (*ũ*_*L*_) for varying (dimensionless) amplitude of oscillations *A* = *Ã/H*. (b) Lagrangian mean velocity profiles ***ũ***_*L*_ (m/s) for (i) *A* = 10^−3^, (ii) *A* = 4.5 · 10^−3^, and (iii) *A* = 5 · 10^−2^. The colours represent the velocity magnitude, and the arrows are velocity vectors. Here, *α* = 4.5, *ε* = 0.003.

For small values of *A, ũ*_*pd*_ is the dominant contribution to *ũ*_*L*_, which can also be seen from figure 5b case (i). At intermediate values of *A* ∼ 4.5 · 10^−3^, *ũ*_*s*_ becomes comparable with *ũ*_*pd*_. The resulting *ũ*_*L*_ profile changes shape (figure 5b case (ii)) as the positive flow (in positive *x*-direction) of *ũ*_*s*_ near the walls becomes comparable to the negative flow of *ũ*_*pd*_. For large values of *A* ∼ 5 · 10^−2^, the steady streaming becomes the dominant component of ***ũ***_*L*_ (figure 5b case (iii)).

### 3b Transport

In section 2a we found the Péclet number *Pe* using a timescale analysis to identify the transport regimes and thereby the appropriate methodology for simplifying the system. We now consider the effective Péclet number characterising solute transport governed by the time-averaged transport equation (10). In this equation, solutes are transported longitudinally by advection in the form of *A*_0_ · *u*_*L*_ and in the transverse direction by a combination of advection and diffusion. As Stokes drift is negligible, the advective velocity can be approximated as ***u***_*L*_ = ***u***_*s*_ + ***u***_*pd*_, separating steady-streaming effects from production–drainage effects. The Péclet number of the time-averaged transport equation *Pe*_*u*_ is therefore decomposed into the steady-streaming Péclet number *Pe*_*s*_ and the production–drainage Péclet number *Pe*_*pd*_, such that *Pe*_*u*_ = *Pe*_*s*_ + *Pe*_*pd*_.

From equation (10), the dimensionless timescales of longitudinal advection and transverse diffusion are *T*_adv_ = 1*/A*_0_*u*_*L*_ and *T*_diff_ = *α*^2^*σ*, respectively. Considering the advective contribution from the steady streaming alone, 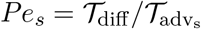 is given by

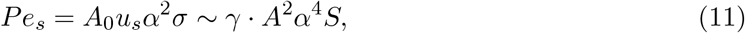

since *u*_*s*_ scales with *Aα*^2^*/ε* with a constant of proportionality *γ* (see section S6). The steady-streaming transport is therefore governed by the amplitude of the oscillations *A*, the frequency of the oscillations through the Womersley number *α*, and the diffusivity of the solutes by the Schmidt number *S*. In particular, fast oscillations, or those with large amplitudes, will result in higher values of *Pe*_*s*_.

The production–drainage transport dynamics are characterised by 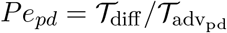

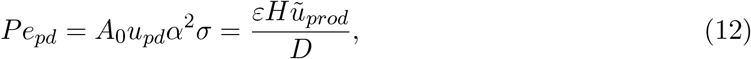

where the final dimensional form clarifies that *Pe*_*pd*_, and therefore transport by ***u***_*pd*_, does not depend on the oscillation parameters *A* and *α*.

Figure 6 shows the variation in *Pe*_*s*_ as the amplitude and frequency of the oscillations vary through *A* and *α*. In the parameter regimes associated with human pulsations, *Pe*_*s*_ is smaller than *Pe*_*pd*_ for the respiratory cycle, but may be comparable for the cardiac cycle and slow waves during sleep. Therefore, we first consider the transport mechanisms that arise for different values of *Pe*_*s*_, before considering the effect of combined contributions of ***u***_*s*_ and ***u***_*pd*_ on solute clearance in the cSAS.

**Figure 6.**
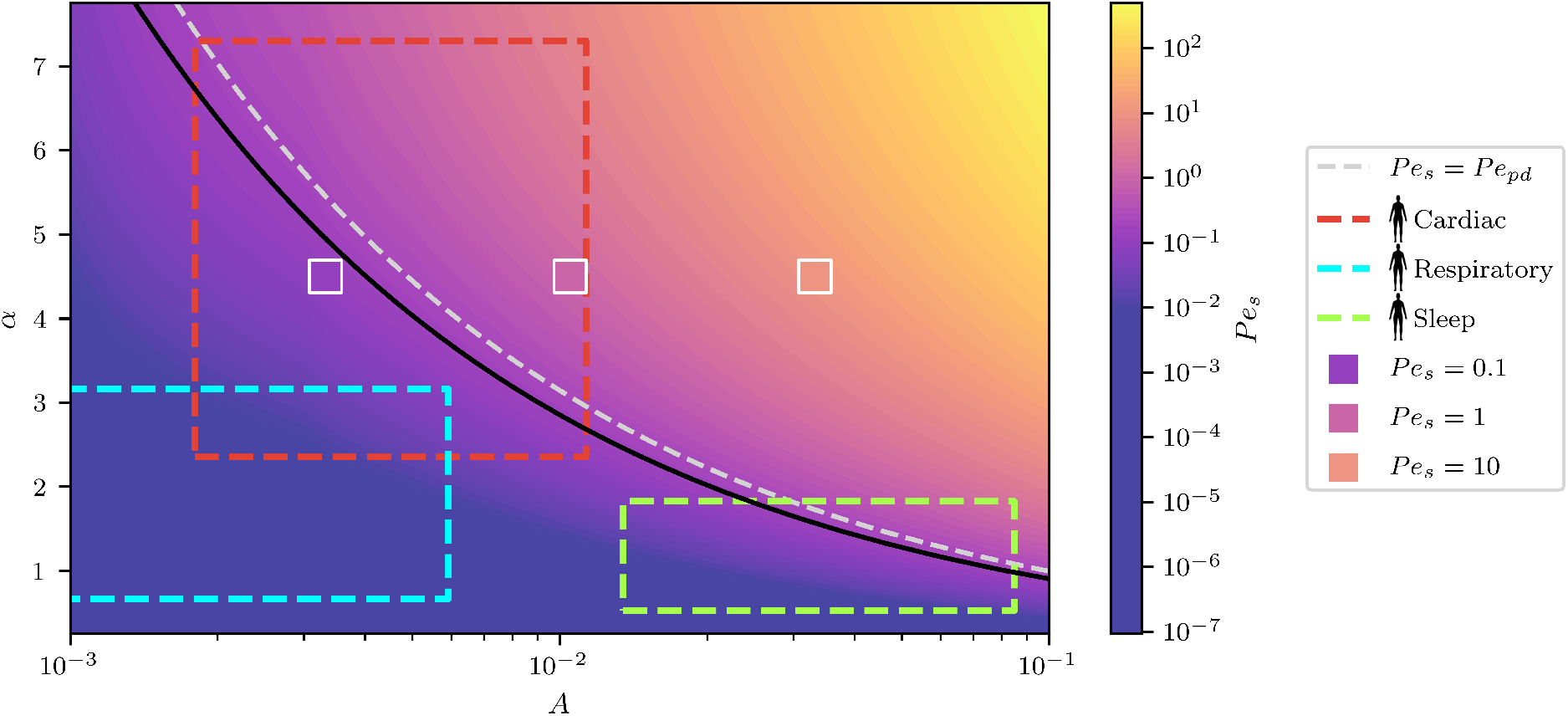
Variation in *Pe*_*s*_ (colourscale) over a range of [*A, α*] values. Contours of constant colour correspond to parameter combinations yielding the same *Pe*_*s*_. The black line indicates transport regime (v) from section 2a, where *Pe* = *α*^2^*AS* = 1*/ε*, i.e. the boundary around and above which we are in transport regime (v). The human cardiac, respiratory and sleep regimes are outlined in red, blue and green boxes, respectively. Most of these physiological cases are within our transport regime. The grey dashed line shows where the advective contribution from ***u***_*pd*_ is equivalent to the contribution from oscillatory flows. The markers at different *Pe*_*s*_ values indicate the values of [*A, α*] used in the simulations in figures 8, 9 and 11. For calculating *Pe*_*s*_ we used *S* = 4096, *ε* = 0.003, *γ* = *ε*max {*u*_*L*_*}/*(*Aα*^2^), and for calculating *Pe*_*pd*_ = 0.21, *D* = *ν/S*, with *ν* from table 1, *ũ*_*prod*_ from table 1 and *H* = 1.45 mm.

In the following sections, we study the transport of solutes in different physiologically motivated scenarios. We consider three test cases for solute transport, schematically shown in figure 7:

**Figure 7.**
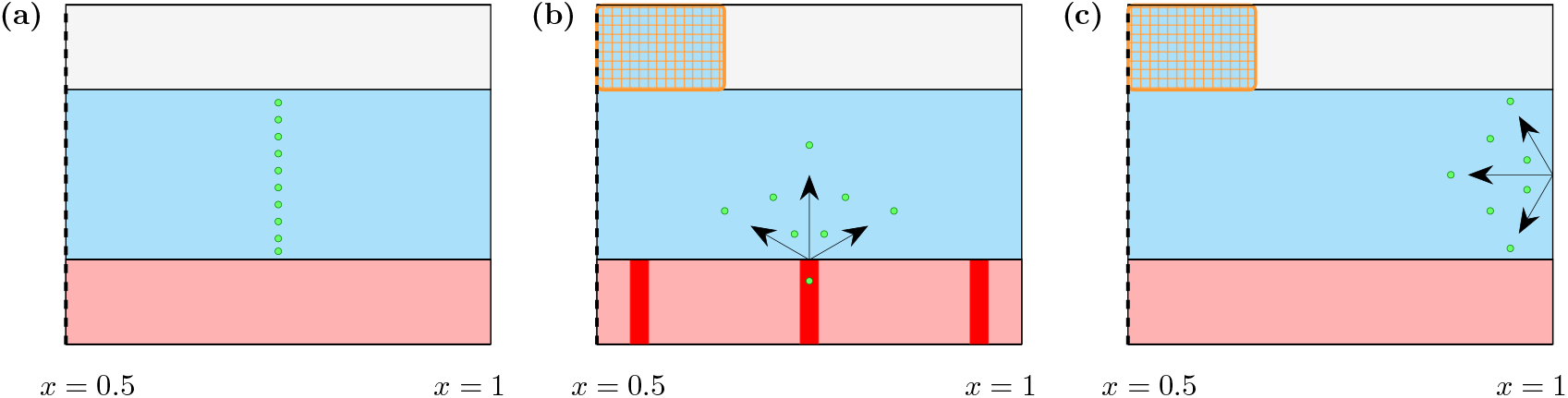
Schematic of physiological case studies. Figure (a) shows local solute initialisation, where the solute is initialised as a Gaussian in the horizontal direction and uniform in the vertical direction at *x* = *x*_0_. Figure (b) shows case (ii), a local source of solute from the brain surface, e.g. from brain tissue or PVS. Figure (c) shows case (iii), an imposed concentration on the spinal boundary, e.g. a solute injected intrathecally in the spinal canal entering the cSAS. In case (ii) and (iii), solute can exit the channel through the intracranial CSF absorption zone, assumed to be arachnoid granulations in human (orange), or by the spinal canal at *x* = 1.

1. A local initialisation of solute in steady-streaming flow (***u***_*pd*_ = 0), where the concentration field is initialised with a Gaussian profile independent of *η*, representing a solute injected into the domain;
2. A solute source from the brain surface representing solute coming from the PVS or from the brain tissue itself, both with and without ***u***_*pd*_;
3. A prescribed solute concentration in the spine, representing the case of intrathecal drug administration.

### 3bi Case study 1: Local initialisation in oscillatory flows

We first explore the case where there is no production–drainage flow, i.e. ***u***_*pd*_ = **0** and the brain and dura membrane are impermeable to the solute, ∂*c/*∂*η* = 0 at *η* = −1, 0, to investigate a solute transported by ***u***_*s*_ only. We prescribe the initial condition 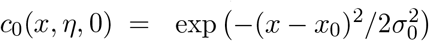, where the Gaussian is centred at *x*_0_ = 0.75 and has standard deviation *σ*_0_ = 0.01. At *x* = 1 we impose a mixed boundary condition for the solute such that for an inward fluid velocity, *u*_*L*_ *<* 0, there is no solute in the fluid (*c* = 0) and for *u*_*L*_ *>* 0, we impose *dc/dx* = 0, which implies that solute leaves the channel with the flow. The latter condition assumes a well-mixed compartment of infinite CSF connected to each end of the channel. The equivalent boundary condition is imposed at *x* = 0.

Figure 8 shows the transport of a solute, initialised as a bolus injection, by ***u***_*s*_ for different *Pe*_*s*_. Figure 8a corresponds to *Pe*_*s*_ = 0.1, representing a regime where transversal diffusion dominates advective forces. This regime does not exhibit effective longitudinal transport.

**Figure 8.**
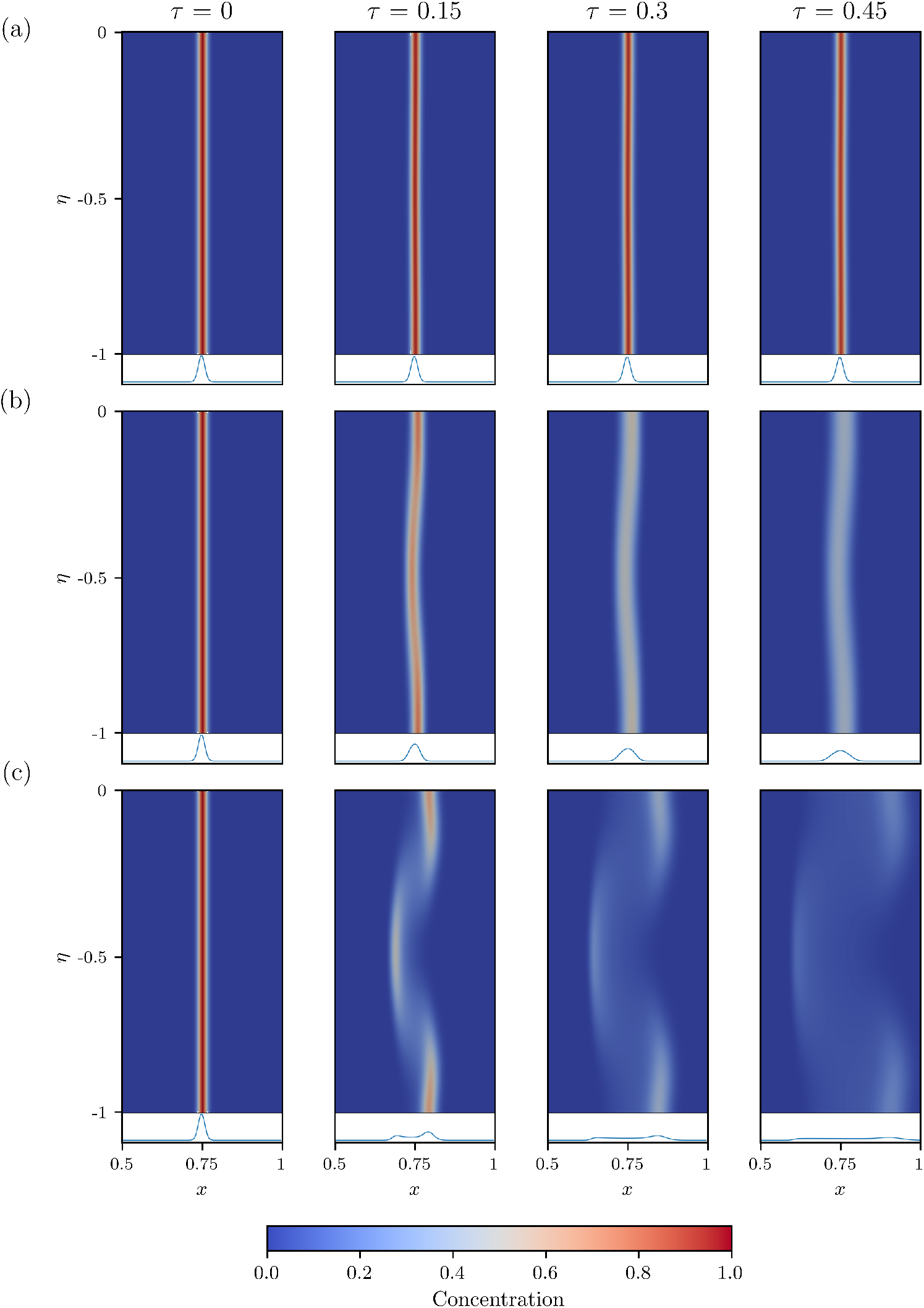
Time snaps of concentration profiles (colourscale) of case study 1 (local solute initialisation in the cSAS) for different *Pe*_*s*_: (a) *Pe*_*s*_ = 0.1, (b) *Pe*_*s*_ = 1 and (c) *Pe*_*s*_ = 10. *Pe*_*s*_ is varied by changing the dimensionless amplitude of oscillations *A*: (a) *A* = 0.00332, (b) *A* = 0.0105, and (c) *A* = 0.0332. The columns represent different (dimensionless) times, as indicated by the titles above the first row. The domain here is the right half of the dimensionless domain: *x* ∈ [1*/*2, 1], *η* ∈ [−1, 0]. The curves under the domains in each panel represent depth-integrated concentration profiles. Here *S* = 4096, *α* = 4.5, *ε* = 0.003 and *γ* = 0.0053.

In the regime shown in figure 8b, with *Pe*_*s*_ = 1, advection balances diffusion. Longitudinal transport is more efficient due to the interplay between transverse diffusion and advection. High transverse diffusion, relative to advection, means the profile is mostly homogeneous in *η*.

Finally, figure 8c shows the advection-dominant regime with *Pe*_*s*_ = 10. In this case, advection overwhelms diffusion and the solute follows the steady-streaming flow profile, diffusing transversely on a slower time scale. This figure highlights the influence of flow on concentration profiles. We note that for high *Pe*_*s*_ concentration profiles are not homogeneous in *η*.

### 3bii Case study 2: Local source from the brain surface

We now consider a solute coming from the surface of the brain. We investigate two subcases: case 2a, a local source, analogous to a leak from PVS or from a small part of the brain, and case 2b where we impose a uniform flux along the entire brain surface, similar to waste product being expelled from the brain. In case 2a, we implement the source as a flux condition in a small region on the lower boundary of the channel, so that 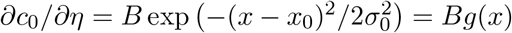 centred at *x*_0_ with *σ*_0_ = 0.01. *B* is chosen to impose a fixed flux of magnitude 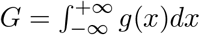, such that 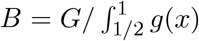. In case 2b, the flux along the lower boundary is constant, and the condition becomes ∂*c*_0_*/*∂*η* = *G*, which ensures the total flux entering the domain is the same in both cases. We prescribe zero concentration in the domain initially, and the same boundary condition as in section 3bi at *x* = 0, 1.

CSF and solutes can clear into the intracranial absorption site, away from the spinal canal, which is modelled by the Kadem-Katchalsky membrane condition equation (S20) ([23]), which at leading order is

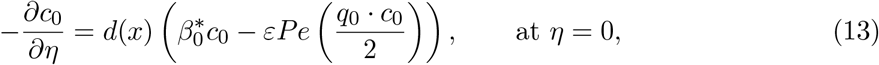

where *d*(*x*) (equation (7)), prescribes the area permeable to both CSF and the solute (sketched in the schematic in figure 10), 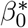 is the dimensionless membrane permeability, and *q*_0_ is the magnitude of dimensionless outlet velocity. This boundary condition assumes that both advection by CSF and solute diffusivity through the membrane contribute to transport across the membrane. For cases 2 and 3, we chose parameters representative of human arachnoid granulations, including the characteristic drainage-region size appearing in *d*(*x*) and the membrane permeability *β*_0_^∗^ (see sections S10(c).3 and S10(d).2). To consider another drainage site, the parameters should be adjusted accordingly. We also assume that solute may exit to the spinal canal, where it can be cleared by lymphatic or venous routes [36].

### Case 2 with oscillatory flow only, *u*_*pd*_ = 0

Figure 9(a, c, e) shows the time-dependent and steady-state profiles of a point source at *x*_0_ = 0.8 for ***u***_*s*_ alone (***u***_*pd*_ = 0). For *Pe*_*s*_ = 0.1, in figure 9a, the solute diffuses vertically across the channel before reaching the dura membrane. The solute permeability of the dura membrane is very small around *x* = 0.8 (see the schematic in figure 10) causing an accumulation of solute concentration. Although the solute spreads to the right over a long period of time (see figure 9a steady state), weak advection leads to little clearance into the spinal canal.

**Figure 9.**
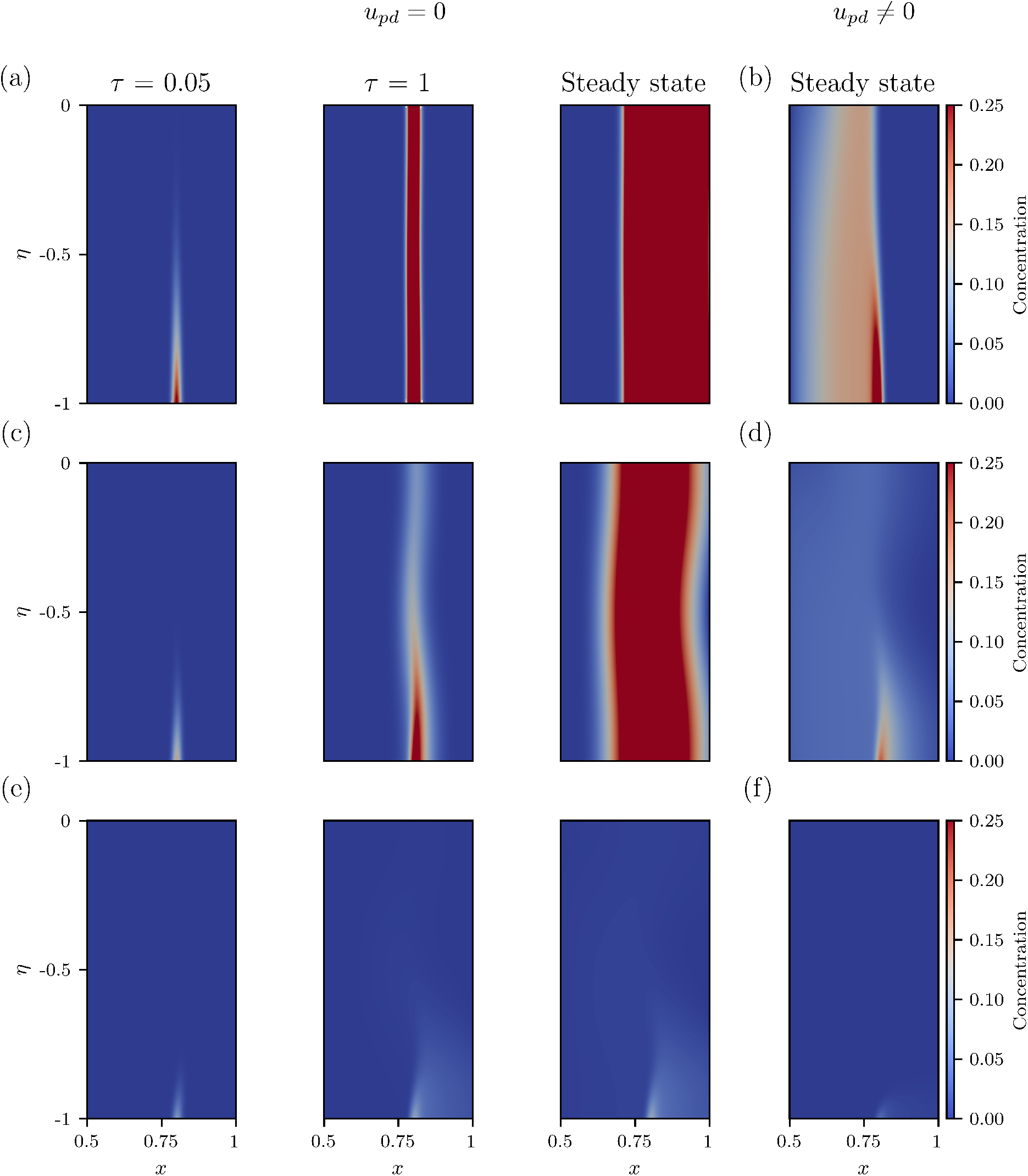
Time snaps and steady states of concentration profiles (colourscale) in the case of point source simulations (case 2a) for different *Pe*_*s*_ : (a,b) *Pe*_*s*_ = 0.1, (c,d) *Pe*_*s*_ = 1 and (e,f) *Pe*_*s*_ = 10. *Pe*_*s*_ is varied by changing the dimensionless amplitude of the oscillations *A*: (a,b) *A* = 0.00332, (c,d) *A* = 0.0105, and (e,f) *A* = 0.0332. The first two columns represent the concentration profile transported by steady streaming only (*u*_*pd*_ = 0) at different (dimensionless) times with the third column showing the resulting steady state, as indicated by the titles above the first row. The final column contains the corresponding steady states when production–drainage flow is included (*u*_*pd*_≠ 0). The domain is the right half of the dimensionless domain: *x* ∈ [1*/*2, 1], *η* ∈ [−1, 0]. Here *S* = 4096, *α* = 4.5, *ε* = 0.003 and *γ* = 0.0053. The source is placed at *x*_0_ = 0.8.

For balanced advection and diffusion (figure 9c), the steady-state concentration profile is vertically heterogeneous, with a higher concentration near the source. Increased advection contributes to greater longitudinal spreading, which, in tandem with vertical diffusion, enhances solute dispersal throughout the channel. This, in turn, enables clearance by both intracranial and spinal pathways.

Finally, in the advection-dominant case (figure 9e), the solute is advected predominantly towards the spine by the positive velocity along the bottom of the channel, where it clears through the spinal canal. There is minimal cross-channel diffusion, and much less of the solute reaches the dura membrane, thus limiting clearance by arachnoid granulations.

To understand the relative importance of cranial absorption sites and spinal canal in solute clearance, we examine the solute mass accumulation in the steady state (figure 10a) and the percentage of solute flux cleared through arachnoid granulations (figure 10b) as *Pe*_*s*_ is varied. At low and intermediate *Pe*_*s*_, total mass increases for sources located closer to the the spine (*x*_0_ = 0.8, green line). This occurs because transverse diffusion favours clearance through the centrally located arachnoid granulations; for sources placed peripherally (*x*_0_ ≳ 0.75), solutes do not reach the drainage site (see figure 10 top panel).

**Figure 10.**
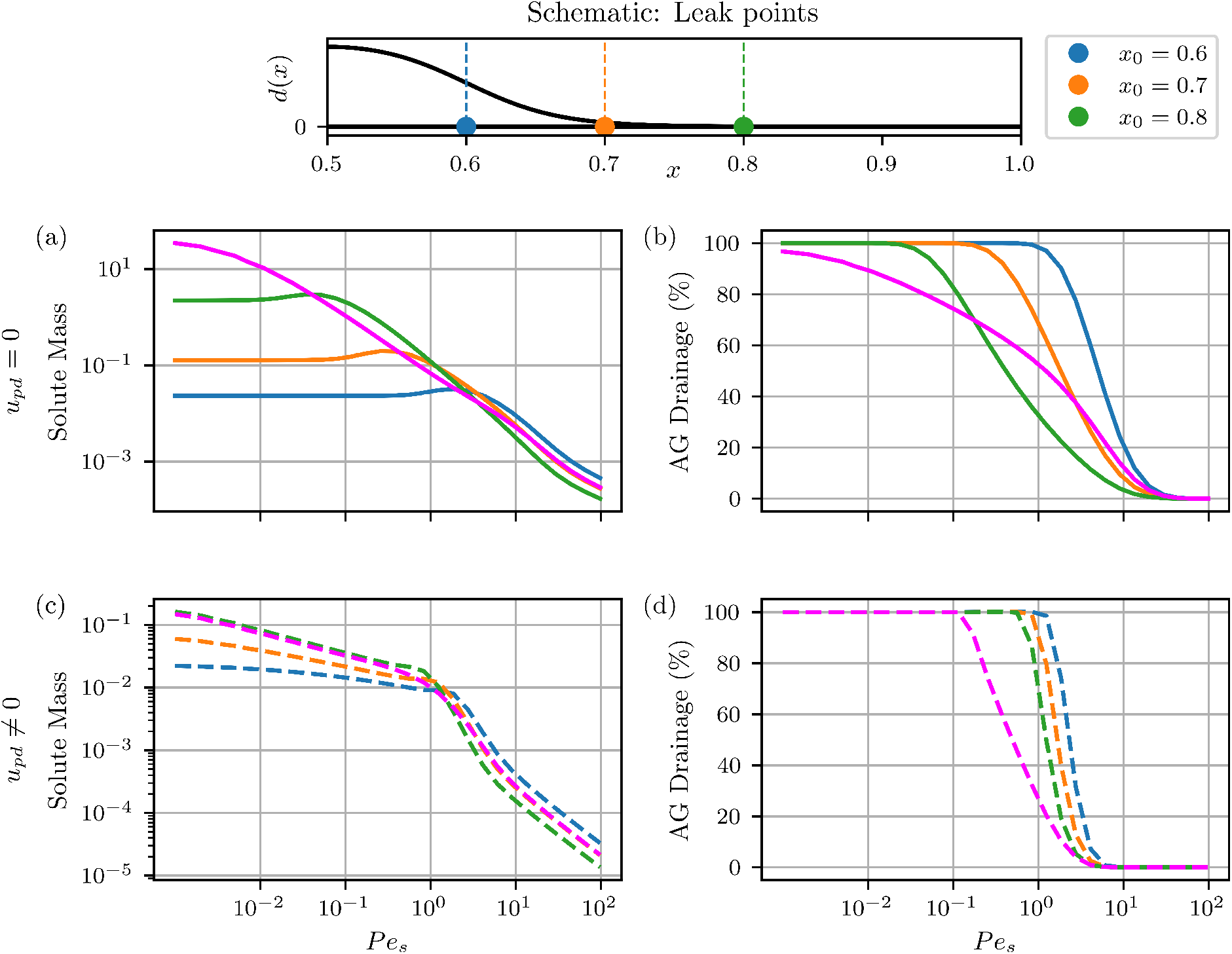
Comparison of mass accumulation and tendency to drain via arachnoid granulations for steady streaming only (*u*_*pd*_ = 0) - solid lines - (a,b), and steady-streaming flows combined with production–drainage flows *u*_*pd*_ ≠ 0-dashed lines - (c,d) *Pe*_*s*_ varied by increasing *A*. The schematic on top shows the function describing the drainage through arachnoid granulations *d*(*x*), and the source sites at *x* = 0.6 (blue), *x* = 0.7 (orange), *x* = 0.8 (green). (a,c) Show the total solute mass accumulated at steady state in the channel from a point source for each source site shown in the schematic with corresponding colours, and for a source along the full brain surface (magenta). (b,d) Show the percentage drainage at steady state through the arachnoid granulations for the same source sites. Note that we use the value of *u*_*pd*_ corresponding to humans in table 1.

For *x*_0_ = 0.8 and low *Pe*_*s*_, the solute is cleared exclusively through arachnoid granulations (figure 10b). As *Pe*_*s*_ increases towards *Pe*_*s*_ ∼ *O*(10^−1^), the solutes are advected away from arachnoid granulations, as we see in figure 9a, while not quite reaching the spine, resulting in an increase of the total mass. When *Pe*_*s*_ is increased further, advection towards the spine strengthens, promoting clearance into the spine, and the total mass decreases.

For source locations closer to the centre of the domain, *x*_0_ = 0.7 (orange) and *x*_0_ = 0.6 (blue), the behaviour is qualitatively similar, except that the peak mass occurs at higher *Pe*_*s*_ values in figure 10a. These sources face the arachnoid granulations as seen in the schematic in figure 10, leading to more efficient clearance by transversal diffusion. Moreover, the longitudinal fluid velocity increases linearly in *x*, being at its weakest at the domain centre (*x* = 0.5) and strongest at the spinal outlet (*x* = 1). Therefore, solutes emitted near the centre experience weaker initial advective transport and must be advected further to reach the spine, requiring higher *Pe*_*s*_ to initiate spinal clearance as seen in figure 10b.

We now consider case 2b, where the source is uniform at the surface of the brain. The steady-state concentration profiles are shown in figure S9. For small *Pe*_*s*_, solute accumulates in the region near the spine (figure 10), where there is weak advection and no clearance via arachnoid granulations. As *Pe*_*s*_ increases, more solute clears into the spine, and the mass decreases.

### Case 2 with production–drainage of CSF, *u*_*pd*_ ≠ 0

In figure 9(b, d, f) we revisit the point source at *x*_0_ = 0.8, with the addition of production– drainage flow. The strength of ***u***_*s*_ is varied by increasing *A* in *Pe*_*s*_, while ***u***_*pd*_ is kept fixed using *q*(*x*) defined for humans in section S10(c). In the case of small *Pe*_*s*_ = 0.1 (figure 9b), solute accumulates towards the center of the channel. In this regime, ***u***_*s*_ is negligible, and the production–drainage flow advects solute towards the arachnoid granulations. Transverse diffusion contributes to clearance through the intracranial absorption site, particularly close to *x* = 0.5.

As *Pe*_*s*_ increases to 1 (figure 9d), ***u***_*s*_ is comparable with both diffusion and ***u***_*pd*_. The steady-streaming flow counteracts the ***u***_*pd*_ close to the upper and lower boundaries of the channel, while slightly enhancing flow along the centre of the channel. Consequently, transverse diffusion combines with strong advection along the centre of the channel to increase cross-channel dispersal for *x < x*_0_. The combination of advection towards the intracranial drainage site by production– drainage flow, and towards the spinal canal by steady streaming leads to solute clearance through both routes, and therefore less solute accumulating in the channel.

For the advection-dominant transport regime (*Pe*_*s*_ = 10) in figure 9f, ***u***_*s*_ dominates ***u***_*pd*_ and transversal diffusion. Solute is therefore primarily cleared through the spinal canal. However, in comparison to the steady state concentration profile in figure 9e, there is enhanced clearance of solute reaching the middle of the channel *η* = − 0.5, where production–drainage flow increases advection towards the intracranial drainage site.

The inclusion of ***u***_*pd*_ results in a substantial reduction in solute accumulation across all *Pe*_*s*_ in figure 10c. Furthermore, the transition from clearance predominantly through intracranial drainage site to spinal canal occurs over a narrower range of *Pe*_*s*_. Production–drainage flow enhances transport towards the intracranial drainage site, particularly for sources located further from the arachnoid granulations. Consequently, the bias towards arachnoid granulation clearance persists to larger values of *Pe*_*s*_ than in the absence of ***u***_*pd*_, while the differences in clearance behaviour between source locations are diminished.

### 3biii Case study 3: Imposed concentration on the spinal boundary

Our third case study is motivated by the injection of drugs intrathecally into the spinal canal. Many therapeutically relevant drugs exhibit limited or variable permeability across the blood– brain barrier; intrathecal delivery provides an alternative route for these drugs to reach the brain directly via the CSF [9]. There is little evidence indicating the quantity of drug reaching the surface of the brain, especially in different brain regions. In this case study, we investigate how far into the cSAS the solute penetrates, and the degree to which the mobility of the solute, and the concentration reaching the surface of the brain is affected by variations is brain pulsations. We perform simulations with and without ***u***_*pd*_ to quantify its potential role in drug delivery to the brain.

We model solute influx into the cSAS following injection into the spinal canal as a constant concentration entering the domain from outside the channel. Specifically, we prescribe a Dirichlet boundary condition *c*_0_ = 1 along the boundaries at *x* = 1 where the longitudinal velocity satisfies *u*_*L*_(1, *η*) *<* 0. A homogeneous Neumann condition, ∂*c*_0_*/*∂*n* = 0, is applied to the remainder of the boundary. The equivalent inflow/outflow condition is applied at *x* = 0. The boundary condition at the dura (*η* = 0) is given by the membrane condition equation (13), while the brain is impermeable to the solute 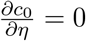 at *η* = −1. We prescribe *c*_0_ = 0 in the domain as the initial condition.

### Case 3 with oscillatory flow only, *u*_*pd*_ = 0

We first examine the transport driven solely by ***u***_*s*_. Steady-state concentration profiles for varying *Pe*_*s*_ (achieved by increasing *A*) are shown in figure 11a. For small *Pe*_*s*_, diffusion dominates and the solute remains in proximity to the spinal outlet. As *Pe*_*s*_ increases, advective transport becomes more effective, pushing the solute further into the domain and enabling some clearance through the arachnoid granulations. However, since ***u***_*s*_ is strongest near *x* = 1 (*x* = 0), the spinal outlet remains the dominant clearance pathway. The fluid velocity is small around *x* = 0.5, which reduces solute advection in this region even at high *Pe*_*s*_.

**Figure 11.**
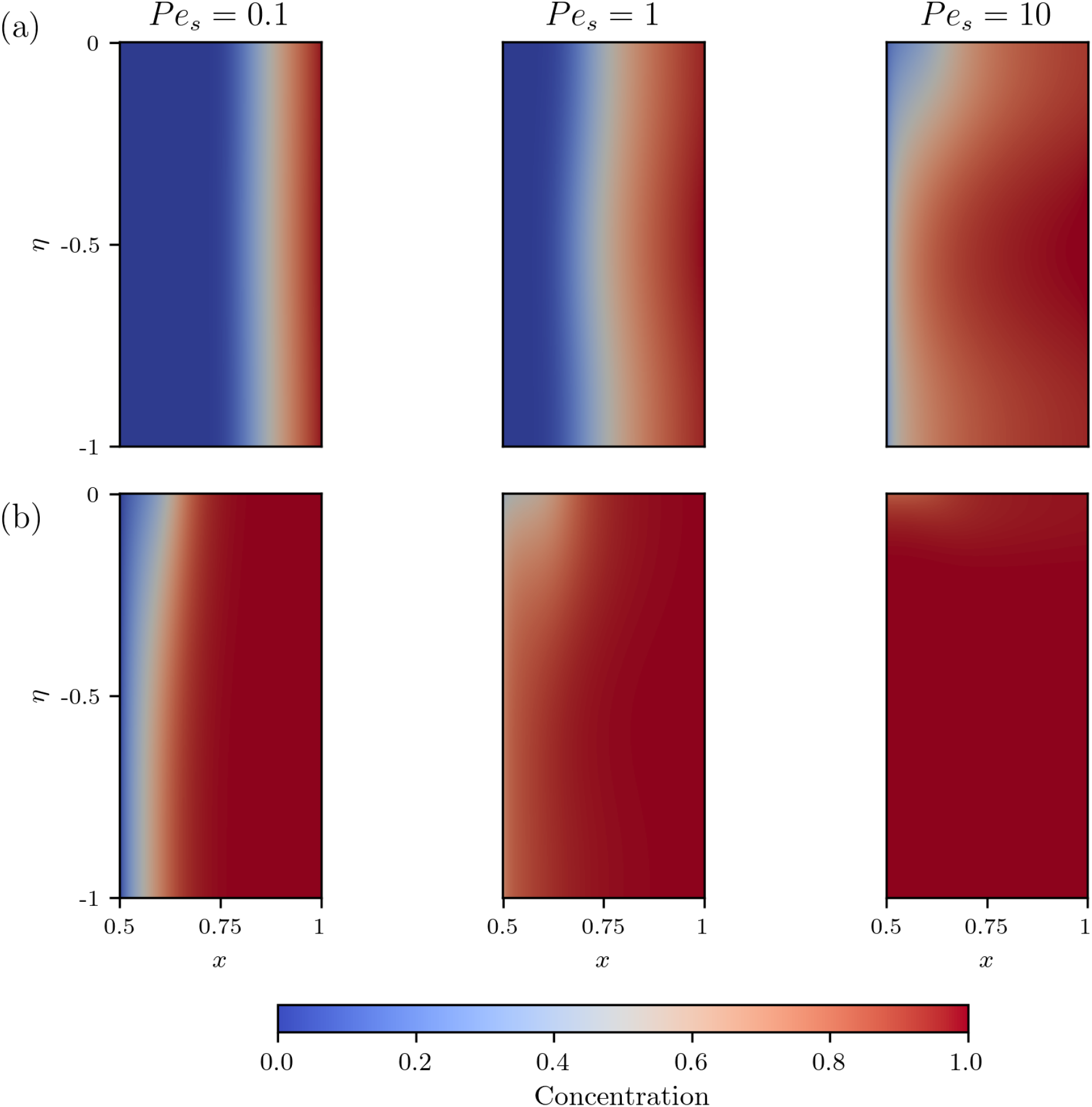
Steady states in the case of a solute coming from the spinal canal for different *Pe*_*s*_ for (a) steady-streaming flow only and (b) ***u***_*pd*_ included. The solute is initialised as *c* = 1 where *u*_*L*_ *<* 0 at the entry to the spinal canal *x* = 1 to mimic a drug delivered intrathecally. The solute is transported further for large *Pe*_*s*_, which is enhanced by the inclusion of ***u***_*pd*_. Here *S* = 4096, *α* = 4.5, *ε* = 0.003, *γ* = 0.0053 and *A* is varied as in the previous case studies.

### Case 3 with production–drainage of CSF, *u*_*pd*_ ≠ 0

Figure 11b shows the corresponding steady states when ***u***_*pd*_ is included. For low and intermediate *Pe*_*s*_ values, ***u***_*pd*_ transports solute into the domain, promoting clearance via the arachnoid granulations and enhancing solute distribution. In comparison with figure 11a, ***u***_*pd*_ increases the depth reached along the brain surface by the front of the concentration profile, with solute extending to the centre (*x* = 0.5) for intermediate *Pe*_*s*_. At high *Pe*_*s*_ steady-streaming advection dominates and solute clearance favours the spinal route. Still, the inclusion of ***u***_*pd*_ facilitates solute transport to the entire brain surface, and enhances clearance through arachnoid granulations.

## 4 Discussion

We investigated the physiological mechanisms by which CSF transports solutes in a simplified 2-dimensional geometry of the cSAS, with emphasis on the effect of previously overlooked steady CSF flows generated by brain pulsations associated with the cardiac cycle, respiration, and vasomotion during sleep. We focused on the role of such steady flows in clearance of metabolic waste from, and drug delivery to, the brain.

Exploiting the small aspect ratio of the domain *ε* and small relative oscillation amplitudes *A*, we defined Péclet numbers *Pe* for the oscillatory flow driven by brain pulsations and identified distinct transport regimes (i)-(v). In our analysis, we focus on regime (v) which is associated with pulsations produced by human physiology, wherein the timescales of *O*(*ε*) steady secondary flows and transverse diffusion are comparable. The physiological parameters associated with the geometry, oscillations, and solute transport are summarised in table 1, with the corresponding parameter estimation detailed in Supplementary Material section S10. Using asymptotic expansions and the separation of two intrinsic timescales, *ω*^−1^ for oscillations and (*εAω*)^−1^ for net solute transport, we derived the Lagrangian mean velocity and corresponding time-averaged transport equation. Net solute transport in the cSAS is mediated by the balance between advection by the Lagrangian mean velocity and transverse diffusion.

The Lagrangian mean velocity consists of three components. The first is the steady streaming, which arises from inertial terms at *O*(*ε*). The steady streaming recirculates CSF; near the brain surface and dura, the flow is directed toward the spine; in the middle of the channel, it moves in reverse toward the domain centre. The second component is production–drainage flow, which is also *O*(*ε*) smaller than the oscillatory flow. Here, we assume that the drainage occurs around the centre of the dura membrane, representing the outflow through an intracranial absorption site. The third term is the Stokes drift, which comes from the interaction between the oscillatory displacement of a fluid particle and the spatial gradients in the oscillatory flow. The relative magnitudes of these three components govern the Lagrangian mean velocity profile.

Our analysis shows that the velocity of production–drainage flow is comparable to the maximum velocity in steady streaming for physiological parameter regimes, implying that both mechanisms jointly govern solute advection. Importantly, production–drainage is constant in magnitude, whereas steady streaming scales quadratically with oscillation frequency and amplitude, becoming the dominant advective term at sufficiently large values of either parameter. The steady streaming is consistently an order of magnitude larger than Stokes drift in the relevant parameter range, rendering the Stokes drift contribution to advection negligible in this simplified description of CSF in the cSAS.

The relative contribution of advective and diffusive terms in the long-time averaged transport equation is captured by the effective oscillatory Péclet number *Pe*_*s*_ = *A*^2^*α*^4^*S*, which depends on (dimensionless) amplitude of oscillations *A*, Womersley number *α* and Schmidt number *S. Pe*_*s*_ determines the importance of steady streaming for solute transport. In humans, for cardiac oscillations and slow waves with large amplitudes, steady streaming plays an important role in solute transport, while it seems to be less relevant for respiratory motion.

To examine transport across parameter regimes, we first considered local initialisation of a solute (case 1) for low, intermediate, and high steady-streaming Péclet number *Pe*_*s*_. For intermediate and large *Pe*_*s*_, the concentration profiles are not transversely homogeneous across the channel, which prevents a description of transport using an effective diffusion coefficient, as previously adapted for transport in mouse PVS [8]. *Pe*_*s*_, which spans a wide physiologically relevant range, governs both the concentration profiles and solute transport rates. This suggests that, to reliably determine transport mechanisms in contrast-agent studies [35], subject-specific measurements of amplitude and frequency of oscillations, as well as cSAS thickness, are required.

We also explored solute transport in two physiologically motivated cases: the clearance of a solute from brain tissue (case 2), and the delivery of a drug injected intrathecally to the brain tissue (case 3). In case 2, we identify several solute accumulation regimes, which in this case are linked with clearance efficiency: i.e., larger solute mass in the domain corresponds to a less efficient clearance. There is a peak in total mass for *Pe*_*s*_ ≈ 0.1–10, suggesting that in this range, the clearance is inefficient. Interestingly, *Pe*_*s*_ for human cardiac and slow-wave pulsations lies in this range (see table 3). We also look at the relative importance of solute fluxes into various absorption routes: for *Pe*_*s*_ *<* 1 the solute is cleared primarily by intracranial absorption, while for *Pe*_*s*_ *>* 10 most solute is cleared into the spinal canal. The inclusion of production–drainage of CSF improves clearance efficiency at lower *Pe*_*s*_, where advection towards the intracranial absorption site enhances solute flux there. This suggests that clearance can be affected by alteration of CSF production in various physiological and pathological conditions.

**Table 3:**
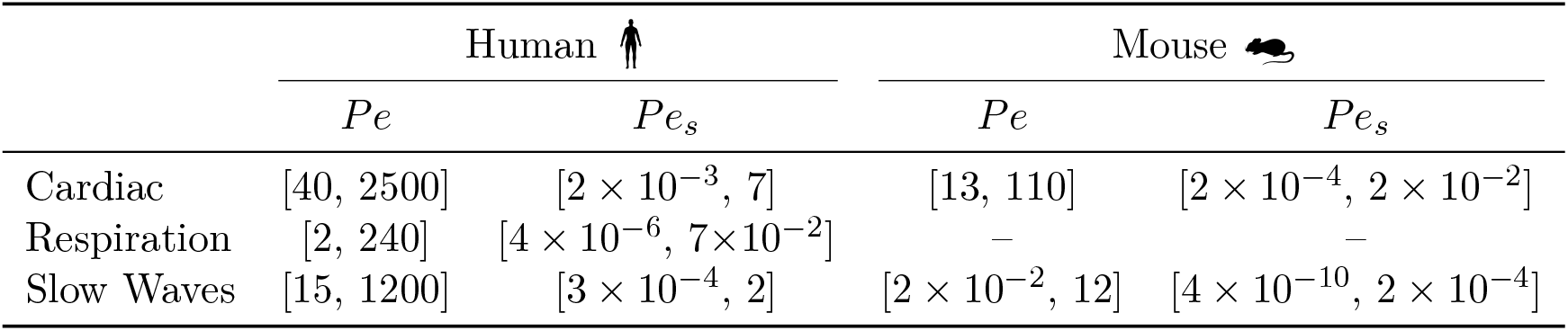
Summary of *Pe* and *Pe*_*s*_ for physiological regimes with *S* = 4096 rounded to 1 or 2 significant figures. In humans *Pe*_*pd*_ = 0.21 and in mice *Pe*_*pd*_ = 2.8 × 10^−4^, calculated from equation (12) with *D* = *ν/S* for *S* = 4096 and the parameter values given in table 1.

|  | Human |  | Mouse |  |
| --- | --- | --- | --- | --- |
| | $Pe$ | $Pe_s$ | $Pe$ | $Pe_s$ |
| Cardiac | [40, 2500] | $[2 \times 10^{-3}, 7]$ | [13, 110] | $[2 \times 10^{-4}, 2 \times 10^{-2}]$ |
| Respiration | [2, 240] | $[4 \times 10^{-6}, 7 \times 10^{-2}]$ | — | — |
| Slow Waves | [15, 1200] | $[3 \times 10^{-4}, 2]$ | $[2 \times 10^{-2}, 12]$ | $[4 \times 10^{-10}, 2 \times 10^{-4}]$ |

In case 3, we found that in the absence of ***u***_*pd*_, a large *Pe*_*s*_ is required to distribute the drug along the domain. Using parameters from figure 11, we estimate that it takes about 9.5 hours for the solute to reach the middle of the domain in the case of ***u***_*pd*_ alone, while with large steady streaming alone (*u*_*pd*_ = 0, *Pe*_*s*_ = 10) it takes around 35 mins. For comparison, solute spreads around the brain in 4–24 hours [41, 35].

Although our model predicts that production–drainage always acts favourably for clearance, the precise drainage pathways remain elusive [36, 40]. Here, we assumed intracranial drainage occurs into the sagittal sinus through arachnoid granulations, as we could approximate parameters for such drainage in humans. However, a recent imaging study did not detect any net flow in cSAS towards the sagittal sinus [55], suggesting that drainage may occur in other sites, such as via the lymphatic vessels in the dorsal dura mater [30, 4]. In mice, solute clearance likely occurs via the cribriform plate into the nasal lymphatics, rather than the arachnoid villi [36]. These alternative intracranial drainage sites, can be incorporated into our model by modifying the drainage area function *d*(*x*) and the parameters associated with ***u***_*pd*_ and membrane permeability.

There is a significant difference in solute transport between mice and humans (figure 2 and table 3). Although cSAS transport in mice also falls into regime (v) for cardiac oscillations, small *Pe*_*s*_ and *Pe*_*pd*_ imply that steady flows have little-to-no effective transport. Vasomotion during sleep lies in regime (iii—iv), corresponding to Taylor-dispersion enhanced transport, which we do not analyse here but has been studied in [8, 28, 27]. The difference is even clearer in smaller spaces such as PVS. For a mouse PVS of thickness ∼ 7 *µ*m and cardiac displacements with *ω* = 2*π* ·10 Hz and *Ã* ≈ 0.12 *µ*m, we estimate *Pe* ≈ 0.27, which corresponds to the Taylor-dispersion regime (iii) (values from [14]). In this case, *α*^2^ = 0.0038 and the steady streaming is negligible. By contrast, for large surface PVS in humans with a width of 0.5 mm [61] and *Ã* = 5 *µ*m (1% displacement of a vessel with 1 mm diameter), we estimate *α*^2^ ≈ 2 and *Pe* ≈ 80, which suggests that steady streaming is non-negligible and the transport is in regime (v). This suggests that, in humans, solute transport is more effective in the cSAS, where steady streaming can contribute to advective transport, whereas in pial PVS small *Pe*_*s*_ implies limited efficacy. In mice, steady streaming is negligible; however, transport in pial PVS lies within a Taylor-dispersion regime, which can enhance effective transport despite weak advection [27]. This interpretation aligns with observations that, in humans, solute spreads around the (healthy) brain in an almost uniform fashion [41], while in mice solute transport is seen primarily along pial PVS rather than in the cSAS [33]. The difference in predicted transport regimes between mice and humans indicates that care should be taken when extrapolating results from one species to another, and when making modelling assumptions.

Both *Pe* and *Pe*_*s*_ are proportional to the Schmidt number *S* = *ν/D*, which is inversely proportional to the diffusion coefficient. Throughout this work, we considered a diffusion coefficient representative of larger solutes, *D* = 1.95 × 10^−10^ m^2^/s, such as amyloid-*β* (*D* ≈ 1.8 × 10^−10^ m^2^/s [58]) or gadobutrol (*D* ≈ 3.8 × 10^−10^ m^2^/s [53]). For solutes with different diffusivities, the relative importance of advection increases as 1*/D*. For example, tau protein, which is central to the pathology of Alzheimer’s disease, has a reported diffusivity of *D* ≈ 6.3 × 10^−11^ m^2^*/*s [42]. This decrease in *D* increases *Pe*_*s*_, implying that advective transport by steady flows becomes more pronounced. For larger tau aggregates, the diffusivity may decrease further, in which case transport is expected to become increasingly dominated by advection by steady flows within the present framework. Conversely, smaller and more diffusive solutes remain in regimes (iv)–(v) for cardiac oscillations in humans and (iii)–(iv) in mice, but with negligible advection by steady streaming, *Pe*_*s*_ ≪ 1.

The simplified model presented here provides insight into the relevant transport regimes in oscillatory flow, highlighting the importance of steady streaming for solute transport in humans. There are, however, several limitations worth mentioning. First, the model relies on the assumption of a small aspect ratio of the cSAS. While this is valid for healthy cSAS, in cases of neurodegeneration, the aspect ratio may become large, and other methods might be considered to study transport.

Second, the model neglects the geometrical complexities of the brain (i.e. shape and sulci). While the rectangular geometry provides a useful idealisation of the cSAS, spatial variations in curvature and thickness are expected to modify the magnitude and direction of steady-streaming flows [26, 2], and hence may alter quantitative predictions of solute transport. The oscillatory and steady-streaming velocity profiles in a spherical and realistic cSAS have been considered in [13]. For an axisymmetric sphere undergoing uniform pulsation, oscillatory and steady-streaming flows are qualitatively the same as in this work. For spatially non-uniform pulsations or more realistic geometries, there may be steady streaming circulation around the brain [13], though this has not been confirmed in imaging studies [54]. Similarly, for asymmetric channels, net flow may be generated [34].

We anticipate that the Péclet number analysis presented here can be extended to identify the dominant transport mechanisms in three-dimensional settings; however, the additional spatial dimension and geometric complexity are expected to modify the characteristic solute transport. Further work is required to clarify both the existence of net flow in the cSAS and the impact of geometrical complexities on solute transport in physiologically realistic geometries.

Third, we model oscillations with a sine wave in time, while real brain surface displacements are spatially inhomogeneous and have more complex temporal dynamics [1]. In section S10, to estimate the amplitudes of oscillations, we assume a spherical brain with spatially uniform displacements and estimate the amplitude from existing MRI measurements of the CSF flows (see table 1) in humans and by inferring volume change from infusion tests or blood vessel wall deformations in mice. This simplification neglects regional heterogeneity, likely underestimating local displacements; human cardiac cycle motion can reach 20-40 *µ*m in certain regions of the brain surface [1], which is larger than our estimate of 6.6 *µ*m. Accounting for geometrical and motion complexities may change the relative importance of steady streaming, and Stokes drift [26]. Furthermore, steady streaming may produce net flow in certain regions, as was observed for the spine [27]. This will also be investigated in future research.

Fourth, the cSAS in our model does not include arachnoid trabeculae, which have been shown to enhance solute transport [43, 49, 57] through dispersion. In the spine, geometry induced mixing, in the form of eddies close to the nerve roots, has been identified as the dominant mechanism behind intrathecal drug dispersion [5, 6], with trabeculae enhancing local solute dispersion [49]. In the cSAS, there are no nerve roots therefore the local dispersion created by trabeculae may be comparable with other advective and diffusive transport mechanisms. Inclusion of trabeculae may also reduce the relative importance of steady streaming for solute transport [34].

Fourth, the cSAS in our model does not include arachnoid trabeculae, which have been shown to enhance solute transport through increased dispersion [43, 49, 57]. In the spinal subarachnoid space, geometry-induced mixing associated with eddies near the nerve roots has been identified as a dominant mechanism driving intrathecal drug dispersion [5, 6], while trabeculae further enhance local solute mixing [49]. In contrast, the cranial subarachnoid space lacks the prominent nerve-root structures present in the spine, suggesting that trabeculae-induced dispersion may play a role comparable to other advective and diffusive transport mechanisms in the cSAS. Additionally, inclusion of trabeculae may reduce the relative importance of steady streaming for solute transport [34]. Inclusion of trabeculae may therefore alter the balance between dispersion and oscillatory transport mechanisms.

Finally, for simplicity, we treat the brain surface as impermeable to solutes. In reality, PVS may provide routes for fluid exchange between the cSAS and brain tissue, though the precise pathways and mechanisms linking these compartments remain debated [24]. Incorporating this effect would require modelling a time-dependent flux across the oscillating brain boundary and the resulting coupling between porous transport and cSAS flow. In the absence of quantitative data characterising this exchange during physiological oscillations, we isolate the transport mechanisms driven by brain motion alone. Allowing for porous exchange would likely modify the magnitude and structure of the steady flows, and hence the resulting solute transport.

## Supporting information

Supplementary Material for derivations and parameter estimation.

## Data accessibility

[GitHub repository](https://github.com/lananeff/2-D-solute-transport-in-cSAS).

## Funding

The authors gratefully acknowledge the financial support of Royal Society International Exchanges award IES\R3\233021 to MD and AV.

## Notes

### Competing Interest Statement

The authors have declared no competing interest.

### Summary of Updates

Updated results for physiologically relevent parameters and other minor corrections.

