## Supplementary Material for derivations and parameter estimation. for "How brain pulsations drive solute transport in the cranial subarachnoid space: insights from a toy model"

[1]School of Mathematics and Maxwell Institute of Mathematical Sciences, University of Edinburgh, Edinburgh, UK
[2]SAINBIOSE INSERM U 1059, École nationale supérieure des Mines de Saint-Étienne, Saint-Étienne, France

#### S1 Summary of the model

In this section, we summarise the equations governing oscillatory fluid flow and solute transport in the 2-dimensional channel $(\tilde{x}, \tilde{y}) \in [0, L] \times [-\tilde{h}(t), 0]$. The fluid motion is given by the 2-dimensional Navier–Stokes equations for an incompressible fluid

$$\rho\left(\frac{\partial \tilde{\boldsymbol{u}}}{\partial \tilde{t}} + (\tilde{\boldsymbol{u}} \cdot \nabla)\tilde{\boldsymbol{u}}\right) = -\nabla \tilde{p} + \mu \nabla^2 \tilde{\boldsymbol{u}}, \tag{S1}$$

$$\nabla \cdot \tilde{\boldsymbol{u}} = 0, \tag{S2}$$

for fluid velocity $\tilde{\boldsymbol{u}} = (\tilde{u}, \tilde{v})$ and pressure $\tilde{p}$, where $\rho$ is the density and $\mu$ is the dynamic viscosity of the fluid. The transport of the solute with concentration $\tilde{c}$ is governed by the advection–diffusion equation

$$\frac{\partial \tilde{c}}{\partial \tilde{t}} + \tilde{\boldsymbol{u}} \cdot \nabla \tilde{c} = \nabla \cdot (D \nabla \tilde{c}), \tag{S3}$$

where $D$ is the diffusion coefficient. The time-periodic oscillation of the lower boundary of the channel is described by $\tilde{h}(\tilde{t}) = H + \tilde{A} \sin(\omega \tilde{t})$.

We impose the no-slip condition at the brain surface, with normal velocity equal to that of the oscillating wall. The dura membrane is a rigid, permeable boundary with prescribed drainage flow $\tilde{q}(x)$ through an intracranial drainage site, and additionally a no-slip condition is imposed on the fluid there. The pressure $\tilde{p} = 0$ is imposed at the ends of the channel to close the fluid problem. The boundary conditions for the fluid problem are summarised in figure S1.

The boundary conditions for the solute are case specific (see section 3). Here we provide more detail on the boundary condition associated with solute absorption into the sagittal sinus

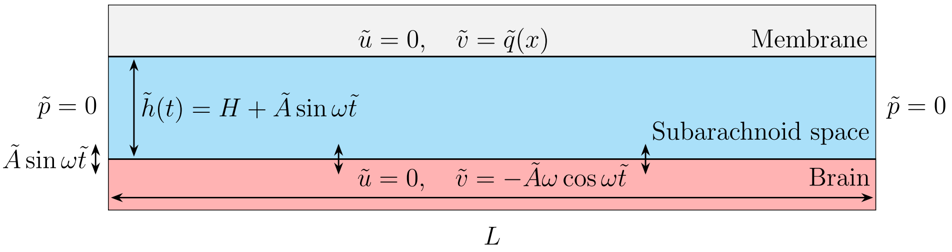


Figure S1: Summary of the simplified fluid-flow problem associated with the flow of CSF in the cSAS driven by pulsations of the brain.

through intracranial drainage sites e.g. arachnoid granulations or the cribriform plate [1]. We model the absorption of the solute using the flux condition

$$\tilde{\boldsymbol{J}} \cdot \boldsymbol{n} = \Phi \quad \text{at} \quad \tilde{y} = \{0, -\tilde{h}(t)\} \quad \text{for} \quad \tilde{\boldsymbol{J}} = -D\nabla\tilde{c} + \tilde{\boldsymbol{u}} \cdot \tilde{c}, \tag{S4}$$

where $\boldsymbol{n}$ is the vector normal to the surface. The function $\Phi(\tilde{x}, \tilde{y}) = 0$ at $\tilde{y} = -\tilde{h}(t)$ where the brain is impermeable to the solute. Where solute transport occurs at the intracranial absorption site at the dura membrane, $\Phi$ is modelled using the Kedem-Katchalsky equation [2] for solute transport across a membrane

$$\Phi(x, y) = \tilde{\beta}(\tilde{c} - \tilde{c}_{ext}) + \tilde{v} \cdot \bar{\tilde{c}}, \tag{S5}$$

where $D$ is the molecular diffusivity of the solute, $\tilde{\beta} = D/H_{AG}$ is the solute permeability, $\tilde{c}_{ext} = 0$ is the concentration of the solute in the sagittal sinus and $\bar{\tilde{c}}$ is the mean solute concentration, assuming $\bar{\tilde{c}} = (\tilde{c} + \tilde{c}_{ext})/2$. Then the dimensional boundary condition for the solute at the intracranial drainage site is

$$-D\frac{\partial \tilde{c}}{\partial \tilde{y}} + \tilde{v} \cdot \tilde{c} = \tilde{\beta}\tilde{c} + \frac{\tilde{v} \cdot \tilde{c}}{2}, \tag{S6}$$

$$-D\frac{\partial \tilde{c}}{\partial \tilde{y}} = \tilde{\beta}\tilde{c} - \frac{\tilde{v} \cdot \tilde{c}}{2}. \tag{S7}$$

#### S2 Non-dimensionalisation

The natural timescale of the problem is the frequency of oscillations such that $\tilde{t} = \omega^{-1}t$. We choose to scale the vertical velocity by the amplitude and frequency of oscillations $\tilde{A}\omega$, so that $V = \tilde{A}\omega$. We use lubrication-type scalings

$$\tilde{x} = Lx, \qquad \tilde{y} = Hy, \qquad \tilde{u} = Uu, \qquad \tilde{v} = Vv, \qquad \tilde{p} = \mu\omega A\varepsilon^{-2}p, \qquad \tilde{A} = AH, \tag{S8}$$

where the dimensional variables are denoted with tildes, the dimensionless parameter $\varepsilon = H/L$ is small. To find the longitudinal velocity scale $U$, we substitute the dimensionless expressions into the continuity equation (S2). The scaling $U = \tilde{A}\omega/\varepsilon = V/\varepsilon$ emerges from the dominant balance. The scale for pressure is chosen to balance the pressure gradient with the dominant viscous term.

The dimensionless equations are

$$\alpha^2 \frac{\partial u}{\partial t} + \varepsilon A_0 \alpha^2 \left( u \frac{\partial u}{\partial x} + v \frac{\partial u}{\partial y} \right) = -\frac{\partial p}{\partial x} + \varepsilon^2 \frac{\partial^2 u}{\partial x^2} + \frac{\partial^2 u}{\partial y^2}, \tag{S9}$$

$$\varepsilon^2 \alpha^2 \left( \frac{\partial v}{\partial t} + \varepsilon A_0 \left( u \frac{\partial v}{\partial x} + v \frac{\partial v}{\partial y} \right) \right) = -\frac{\partial p}{\partial y} + \varepsilon^2 \left( \varepsilon^2 \frac{\partial^2 v}{\partial x^2} + \frac{\partial^2 v}{\partial y^2} \right), \tag{S10}$$

$$\frac{\partial u}{\partial x} + \frac{\partial v}{\partial y} = 0, \tag{S11}$$

$$\frac{\partial c}{\partial t} + \varepsilon A_0 \left( u \frac{\partial c}{\partial x} + v \frac{\partial c}{\partial y} \right) = \frac{1}{\alpha^2 S} \left( \varepsilon^2 \frac{\partial^2 c}{\partial x^2} + \frac{\partial^2 c}{\partial y^2} \right). \tag{S12}$$

The dimensionless parameters which appear in equations (S9) to (S12) are

$$\varepsilon = H/L \sim 10^{-2}, \quad \alpha^2 = \frac{\omega H^2}{\nu} \sim 2\pi \sim O(1), \quad A_0 = \frac{A}{\varepsilon} \sim O(1), \quad S = \frac{\nu}{D} \sim O(\varepsilon^{-2}),$$

including $\alpha$, the Womersley number characterising the relationship between the oscillatory inertia and the kinematic viscosity $\nu$, and $S$, the Schmidt number describing the ratio between $\nu$ and the diffusivity coefficient $D$.

In dimensionless form, the channel spans $(x, y) \in [0, 1] \times [-h(t), 0]$ with $-h(t) = -1 - A\sin(t)$. The boundary conditions for the fluid problem given by equations (S9) to (S11) are

$$u(x, 0) = 0, \qquad v(x, 0) = \varepsilon q(x), \qquad p(0, y) = 0, \tag{S13}$$

$$u(x, -h) = 0, \qquad v(x, -h) = -\cos(t), \qquad p(1, y) = 0, \tag{S14}$$

where the function $q(x)$ derived in section S10(c).3 describes the fluid flow at the arachnoid granulations in humans.

The boundary conditions associated with solute transport (equation (S12)) at $y = 0$ is a homogenous Robin condition

$$-\frac{\partial c}{\partial y} = \beta^* c - A\alpha^2 S \frac{v \cdot c}{2}, \tag{S15}$$

where the value of $\beta^* = \tilde{\beta} H/D$ is approximated in section S10(d).2.

#### S3 Change of variables

We perform a change of variable $\eta = y/h(t)$ to work in the fixed domain by rescaling the transverse coordinate with the deformation of the boundary. In particular, this change of coordinates gives rise to an additional term from the time-derivative

$$\begin{aligned} \frac{\partial}{\partial t} &= \frac{\partial}{\partial x}\frac{\partial x}{\partial t} + \frac{\partial}{\partial \eta}\frac{\partial \eta}{\partial t} + \frac{\partial}{\partial t}\frac{\partial t}{\partial t}, \\ &= -\frac{\eta}{h(t)}\frac{\partial h(t)}{\partial t}\frac{\partial}{\partial \eta} + \frac{\partial}{\partial t}. \end{aligned}$$

Equations (S9) to (S12) in the new coordinate system are

$$\alpha^2\left(\frac{\partial u}{\partial t}+\varepsilon A_0\left(u\frac{\partial u}{\partial x}+\frac{1}{h(t)}v\frac{\partial u}{\partial \eta}\right)-\frac{\eta}{h(t)}\frac{\partial h(t)}{\partial t}\frac{\partial u}{\partial \eta}\right)=-\frac{\partial p}{\partial x}+\varepsilon^2\frac{\partial^2 u}{\partial x^2}+\frac{1}{h^2(t)}\frac{\partial^2 u}{\partial \eta^2}, \tag{S16}$$

$$\varepsilon^2\alpha^2\left(\frac{\partial v}{\partial t}+\varepsilon A_0\left(u\frac{\partial v}{\partial x}+\frac{1}{h(t)}v\frac{\partial v}{\partial \eta}\right)-\frac{\eta}{h(t)}\frac{\partial h(t)}{\partial t}\frac{\partial v}{\partial \eta}\right)=-\frac{1}{h(t)}\frac{\partial p}{\partial \eta}+\varepsilon^2\left(\varepsilon^2\frac{\partial^2 v}{\partial x^2}+\frac{1}{h^2(t)}\frac{\partial^2 v}{\partial \eta^2}\right), \tag{S17}$$

$$\frac{\partial u}{\partial x}+\frac{1}{h(t)}\frac{\partial v}{\partial \eta}=0, \tag{S18}$$

$$\frac{\partial c}{\partial t}+\varepsilon A_0\left(u\frac{\partial c}{\partial x}+\frac{1}{h(t)}v\frac{\partial c}{\partial \eta}\right)-\eta\frac{1}{h(t)}\frac{\partial h(t)}{\partial t}\frac{\partial c}{\partial \eta}=\frac{1}{\alpha^2 S}\left(\varepsilon^2\frac{\partial^2 c}{\partial x^2}+\frac{1}{h^2(t)}\frac{\partial^2 c}{\partial \eta^2}\right). \tag{S19}$$

The boundary conditions must also be transformed to the new coordinate system, where the channel walls are now located at $\eta=-1,0$. For the fluid, the boundary conditions remain the same, though those applied at $y=-h(t)$ are now applied at $\eta=-1$.

The dimensionless boundary condition for the concentration (S15) at $\eta=0$ is

$$-\frac{1}{h(t)}\frac{\partial c}{\partial \eta}=\beta^* c-A\alpha^2 S\left(\frac{v\cdot c}{2}\right). \tag{S20}$$

### S4 Eulerian flow solution

Solutions to the leading-order oscillatory flow and the $O(\varepsilon)$ steady flows can be derived by introducing asymptotic expansions for each of the velocity and pressure variables

$$u=u_0+\varepsilon u_1+O(\varepsilon^2),\quad v=v_0+\varepsilon v_1+O(\varepsilon^2),\quad p=p_0+\varepsilon p_1+O(\varepsilon^2), \tag{S21}$$

and writing the function describing the oscillation of the brain surface as

$$h(t)=1+\varepsilon A_0\sin(t)=1+\varepsilon A_0 h_1(t). \tag{S22}$$

In particular, the derivative $h'(t)=\varepsilon A_0 h_1'(t)$. Since $\varepsilon$ is small, we can Taylor expand the functions $1/h(t)$ and $1/h^2(t)$ expand about $\varepsilon=0$.

#### S4(a) Leading-order oscillatory flow

At leading-order, equations (S16) to (S18) reduce to the linear equations

$$\alpha^2\frac{\partial u_0}{\partial t}=-\frac{\partial p_0}{\partial x}+\frac{\partial^2 u_0}{\partial \eta^2}, \tag{S23}$$

$$0=-\frac{\partial p_0}{\partial \eta}, \tag{S24}$$

$$\frac{\partial u_0}{\partial x}+\frac{\partial v_0}{\partial \eta}=0. \tag{S25}$$

The boundary conditions at leading-order are $u_0, v_0=0$ at $\eta=0$, $u_0=0, v_0=-\cos t$ at $\eta=-1$ and $p_0=0$ at $x=0,1$.

Since the oscillations of the brain surface are harmonic, exact solutions to these equations are sought by resolving $u_0$, $v_0$ and $p_0$ into complex temporal harmonics

$$u_0 = u_{01}e^{it} + \overline{u}_{01}e^{-it}, \qquad v_0 = v_{01}e^{it} + \overline{v}_{01}e^{-it}, \qquad p_0 = p_{01}e^{it} + \overline{p}_{01}e^{-it}.$$

The cosine function can be written in its harmonic form as $\cos t = (e^{it} + e^{-it})/2$.

The solution for the leading-order longitudinal velocity component, in terms of $p_{01}$, is found by expanding equation (S23) in harmonics, integrating twice with respect to $\eta$ and applying the boundary conditions for $u_{01}$ at $\eta = 0, -1$. Then

$$u_{01}(x,\eta) = \frac{1}{\zeta^2}\frac{\partial p_{01}}{\partial x}f(\eta),$$

where

$$f(\eta) = \xi e^{\zeta\eta} + (1-\xi)e^{-\zeta\eta} - 1, \qquad \xi = \frac{e^{\zeta}-1}{e^{\zeta}-e^{-\zeta}}, \qquad \zeta = \sqrt{i\alpha^2}.$$

The expression for leading-order pressure

$$p_{01}(x) = -\frac{\zeta^2}{4J}\left(x^2 - x\right), \qquad J = \frac{1}{\zeta}\left((1-\xi)(e^{\zeta}-1) - \xi(e^{-\zeta}-1) - \zeta\right)$$

is found by integrating (S25) from $\eta = -1, 0$ using the boundary conditions $p_{01} = 0$ at $x = 0, 1$.

Finally the transverse velocity component is derived by integrating equation (S25) once more, this time from $\eta$ to 0. The transverse velocity component is

$$v_{01}(\eta) = -\frac{1}{2J}F(\eta),$$

with

$$F(\eta) = \int_{\eta}^{0} f(\eta)d\eta = \frac{1}{\zeta}\left((1-\xi)(e^{-\zeta\eta}-1) - \xi(e^{\zeta\eta}-1) + \zeta\eta\right),$$

after using the boundary conditions for $v$ to find an explicit expression. Note that due to the direction of integration $F'(\eta) = -f(\eta)$.

The leading-order solution for the oscillatory flow is therefore

$$u_0(x,\eta) = -\frac{1}{2}\left(\frac{2x-1}{2}\right)\frac{f(\eta)}{J}e^{it} + c.c., \qquad v_0(\eta) = -\frac{1}{2}\frac{F(\eta)}{J}e^{it} + c.c..$$

#### S4(b) Steady flows

Time-independent steady streaming is present at $O(\varepsilon)$ due to the nonlinear effects from $(\boldsymbol{u}\cdot\nabla)\boldsymbol{u}$ in section 2.2, and a steady production–drainage flow is also expected on this order from the scaling analysis in section 2.2. The equations governing the steady flow

$$\alpha^2\left\langle 2A_0h_1\frac{\partial u_0}{\partial t}+\frac{\partial u_1}{\partial t}+A_0\left(u_0\frac{\partial u_0}{\partial x}+v_0\frac{\partial u_0}{\partial \eta}-\frac{\partial h_1}{\partial t}\eta\frac{\partial u_0}{\partial \eta}\right)\right\rangle=\left\langle -2A_0h_1\frac{\partial p_0}{\partial x}-\frac{\partial p_1}{\partial x}+\frac{\partial^2 u_1}{\partial \eta^2}\right\rangle, \tag{S26}$$

$$0=\left\langle -\frac{\partial p_1}{\partial \eta}\right\rangle, \tag{S27}$$

$$\left\langle A_0h_1\frac{\partial u_0}{\partial x}+\frac{\partial u_1}{\partial x}+\frac{\partial v_1}{\partial \eta}\right\rangle=0. \tag{S28}$$

consist of the $O(\varepsilon)$ terms in the asymptotic expansion of (S9) - (S11), averaged over the timescale of oscillations $\frac{1}{2\pi}\int_0^{2\pi}\cdot dt=\langle\cdot\rangle$. To solve for the $O(\varepsilon)$ steady flow, we expand the $O(\varepsilon)$ pressure and flow components in harmonics such as

$$\boldsymbol{u}_1=\boldsymbol{u}_{10}+\boldsymbol{u}_{11}e^{it}+\boldsymbol{u}_{12}e^{2it}+c.c.,$$
$$p_1=p_{10}+p_{11}e^{it}+p_{12}e^{2it}+c.c.,$$

where the steady component $\boldsymbol{u}_{10}=\boldsymbol{u}_s+\boldsymbol{u}_{pd}$ is composed of steady streaming and production–drainage flow. Following the harmonic expansion it is clear that the average of the time derivative $\langle u_{1t}\rangle=0$.

Substituting these harmonics and the leading-order solutions into (S26)-(S28) and letting $k=\frac{\alpha^2}{4|J|^2}$ we obtain the system of equations to be solved for steady flows

$$\frac{\partial^2 u_s}{\partial \eta^2}+\frac{\partial^2 u_{pd}}{\partial \eta^2}=kA_0\left(\frac{2x-1}{2}\right)\left(2f\overline{J}+|f|^2+\overline{F}f'+\eta f'\overline{J}+2\overline{J}+c.c.\right)+\frac{\partial p_s}{\partial x}+\frac{\partial p_{pd}}{\partial x}, \tag{S29}$$

$$\frac{\partial p_s}{\partial \eta}+\frac{\partial p_{pd}}{\partial \eta}=0, \tag{S30}$$

$$\frac{\partial u_s}{\partial x}+\frac{\partial u_{pd}}{\partial x}+\frac{\partial v_s}{\partial \eta}+\frac{\partial v_{pd}}{\partial \eta}=i\frac{A_0}{4}\frac{f(\eta)}{J}+c.c.. \tag{S31}$$

The problem can be decomposed into two parts due to the linearity of the system. We first solve the steady streaming problem, which retains the forcing terms arising from oscillatory flow, and then solve for the production–drainage flow.

#### S4(c) Steady streaming

Letting $\boldsymbol{u}_s=\boldsymbol{u}_1^s+\overline{\boldsymbol{u}}_1^s$ and considering separately the forcing terms and their conjugates, the equations governing the steady-streaming problem are

$$\frac{\partial^2 u_1^s}{\partial \eta^2}=A_0\left(\frac{2x-1}{2}\right)\mathcal{F}_1+\frac{\partial p_1^s}{\partial x}, \tag{S32}$$

$$\frac{\partial p_1^s}{\partial \eta}=0, \tag{S33}$$

$$\frac{\partial u_1^s}{\partial x}+\frac{\partial v_1^s}{\partial \eta}=A_0\mathcal{F}_2, \tag{S34}$$

| Steady streaming | Production–drainage |
|---|---|
| $u_s = 0$ at $\eta = -1, 0$ | $u_{pd} = 0$ at $\eta = -1, 0$ |
| $v_s = 0$ at $\eta = -1, 0$ | $v_{pd} = 0$ at $\eta = -1$ |
| | $v_{pd} = q(x)$ at $\eta = 0$ |
| $p_s = 0$ at $x = 0, 1$ | $p_{pd} = 0$ at $x = 1$ |
| | $\frac{\partial p_{pd}}{\partial x} = 0$ at $x = 1/2$ |

Table S1: Summary of boundary conditions associated with the $O(\varepsilon)$ steady steaming and production–drainage flow.

with

$$\mathcal{F}_1 = k\big(2f\overline{J} + |f|^2 + \overline{F}f' + \eta f'\overline{J} + 2\overline{J}\big),$$
$$\mathcal{F}_2 = \frac{i}{4}\frac{f(\eta)}{J}.$$

Integrating equation (S32) twice with respect to $\eta$ such that

$$T(\eta) = \int\int \mathcal{F}_1 d\eta d\eta,$$

and applying the boundary conditions for $u^s$ we obtain the solution

$$u_1^s = A_0\left(\frac{2x-1}{2}\right)(T(\eta) + B_1\eta - T_0) + \frac{1}{2}\frac{\partial p_1^s}{\partial x}(\eta^2 + \eta), \tag{S35}$$

for $u_1^s$ in terms of $p_1^s$, with constants $B_1 = T(-1) - T(0)$ and $T_0 = T(0)$. The expression for $p_1^s$ is obtained by integrating equation (S34) from $\eta = -1$ to $\eta = 0$

$$p_1^s = 6A_0(B_{11} - \frac{i}{4})(x^2 - x), \qquad B_{11} = \int_{-1}^{0}\Big(T(\eta)d\eta + B_1\eta - T_0\Big)d\eta.$$

The transverse steady-streaming component is found by integrating the continuity equation from $\eta$ to $\eta = 0$, and the resulting steady-streaming flow is

$$u_s(x,\eta) = A_0\left(\frac{2x-1}{2}\right)\left(T(\eta) + 6\left(B_{11} - \frac{i}{4}\right)(\eta^2+\eta) + B_1\eta - T_0\right) + c.c., \tag{S36}$$
$$v_s(\eta) = A_0\left(-\frac{i}{4}\frac{F(\eta)}{J} + R(\eta) - 6\left(B_{11} - \frac{i}{4}\right)\left(\frac{\eta^3}{3} + \frac{\eta^2}{2}\right)\right) + c.c., \tag{S37}$$

where $R(\eta) = \int_\eta^0 \Big(T(\eta)d\eta + B_1\eta - T_0\Big)d\eta$.

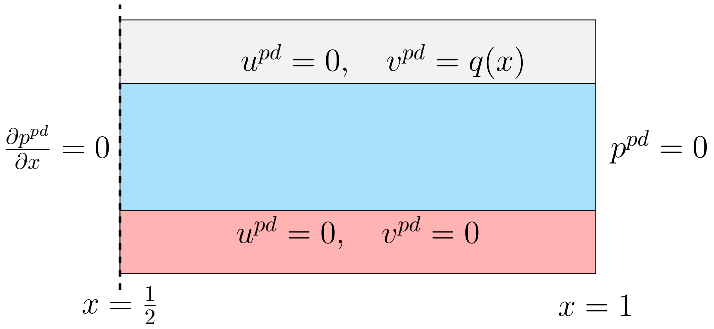


Figure S2: Boundary conditions for production–drainage flow.

#### S4(d) Production–drainage flow derivation

The equations governing production–drainage flow are

$$\frac{\partial^2 u_{pd}}{\partial \eta^2} = \frac{\partial p_{pd}}{\partial x}, \tag{S38}$$

$$\frac{\partial p_{pd}}{\partial \eta} = 0, \tag{S39}$$

$$\frac{\partial u_{pd}}{\partial x} + \frac{\partial v_{pd}}{\partial \eta} = 0. \tag{S40}$$

The boundary conditions associated with production–drainage flow are shown in figure S2 for the half plane. We solve the flow in this plane and use symmetry properties to project this solution across the axis of symmetry. Integrating equation (S38) with respect to $\eta$ and applying the boundary conditions for $u_{pd}$ we obtain,

$$u_{pd} = \frac{1}{2}\frac{\partial p_{pd}}{\partial x}(\eta^2 + \eta). \tag{S41}$$

The pressure component $\frac{\partial p_{pd}}{\partial x}$ is calculated by integrating equation (S40) from $\eta = -1$ to $\eta = 0$, integrating once with respect to $x$ and applying the symmetry condition,

$$\frac{\partial p_{pd}}{\partial x} = 12\big(Q(x) - Q(1/2)\big), \tag{S42}$$

where $Q(x) = \int q(x)dx$. The transverse component $v_{pd}$ is derived by integrating equation (S40) from $\eta$ to $\eta = 0$, and the production–drainage flow is

$$u_{pd}(x,\eta) = 6\big(Q(x) - Q(1/2)\big)(\eta^2 + \eta), \tag{S43}$$

$$v_{pd}(x,\eta) = q(x)\left(1 - 6\left(\frac{\eta^3}{3} + \frac{\eta^2}{2}\right)\right). \tag{S44}$$

#### S5 Lagrangian velocity

The fluid particle trajectories in this flow are influenced by both the steady $O(\varepsilon)$ flows and contributions from the oscillatory flow known as Stokes drift. In order to understand the advective contributions to solute transport in this flow, we consider the particle trajectories, or Lagrangian velocity, of the flow. The displacement of fluid particles in the oscillatory flow occur on two timescales: the timescale of oscillations $\omega^{-1}$ and the timescale of leading order particle transport $\varepsilon^2 A_0 \omega^{-1}$. Particle trajectories in the $(x, \eta)$-domain are defined by

$$\frac{dx_p}{dt} = \varepsilon A_0 u, \qquad \frac{dy_p}{dt} = h(t)\frac{d\eta_p}{dt} + \frac{dh(t)}{dt}\eta_p = \varepsilon A_0 v. \tag{S45}$$

The timescale of the long-time fluid particle trajectories, $\tau = t/(\varepsilon^2 A_0)$, on which relative displacements of $O(1)$ occur can be obtained by performing a dimensional analysis on (S45). The existence of the two timescales in the problem can be exploited to find the particle trajectories in a multiple-timescale expansion

$$\frac{\partial x_p}{\partial t} + \varepsilon^2 A_0 \frac{\partial x_p}{\partial \tau} = \varepsilon A_0 u, \qquad h(t)\left(\frac{\partial \eta_p}{\partial t} + \varepsilon^2 A_0 \frac{\partial \eta_p}{\partial \tau}\right) + \frac{dh(t)}{dt}\eta_p = \varepsilon A_0 v. \tag{S46}$$

We expand the variables $x_p$ and $\eta_p$ asymptotically

$$x_p = x_p^0 + \varepsilon x_p^1 + \varepsilon^2 x_p^2 + O(\varepsilon^3), \qquad \eta_p = \eta_p^0 + \varepsilon \eta_p^1 + \varepsilon^2 \eta_p^2 + O(\varepsilon^3),$$

where each component is assumed to be periodic in $t$. The velocities are Taylor expanded about $\boldsymbol{x_p^0} = (x_p^0, \eta_p^0)$

$$u = u_0(\boldsymbol{x_p^0}, t) + \varepsilon\left(u_1(\boldsymbol{x_p^0}, t) + x_p^1 \frac{\partial u_0}{\partial x}(\boldsymbol{x_p^0}, t) + \eta_p^1 \frac{\partial u_0}{\partial \eta}(\boldsymbol{x_p^0}, t)\right) + O(\varepsilon^2),$$
$$v = v_0(\boldsymbol{x_p^0}, t) + \varepsilon\left(v_1(\boldsymbol{x_p^0}, t) + x_p^1 \frac{\partial v_0}{\partial x}(\boldsymbol{x_p^0}, t) + \eta_p^1 \frac{\partial v_0}{\partial \eta}(\boldsymbol{x_p^0}, t)\right) + O(\varepsilon^2).$$

Substituting these expansions into (S46) writing $h(t) = 1 + Ah_1(t)$ where $A \sim O(\varepsilon)$ once more and gathering terms at each order of $\varepsilon$, we obtain the leading-order equations

$$\frac{\partial x_p^0}{\partial t} = 0, \qquad \frac{\partial \eta_p^0}{\partial t} = 0,$$

which imply that at leading-order the trajectory of the particles only depend on the long-timescale $\tau$.

At $O(\varepsilon)$, the terms are

$$\frac{\partial x_p^1}{\partial t} = A_0 u_0, \qquad \frac{\partial \eta_p^1}{\partial t} + A_0 \frac{dh_1}{dt}\eta_0 = A_0 v_0,$$

which can be integrated with respect to the short time-scale $t$ to give

$$x_p^1 = A_0\left(\int u_0 dt + \hat{x}_p^1\right), \qquad \eta_p^1 = A_0\left(\int v_0 dt - h_1\eta_0 + \hat{\eta}_p^1\right), \tag{S47}$$

where $\hat{\boldsymbol{x}}$ are constants of integration. Expressions for $x_0$ and $\eta_0$ can be recovered at $O(\varepsilon^2)$. We write the $O(\varepsilon^2)$ equations as

$$\frac{\partial x_p^2}{\partial t} + A_0 \frac{dx_p^0}{d\tau} = A_0 \left( u_1 + x_p^1 \frac{\partial u_0}{\partial x} + \eta_p^1 \frac{\partial u_0}{\partial \eta} \right),$$

$$A_0 h_1 \frac{\partial \eta_p^1}{\partial t} + \frac{\partial \eta_p^2}{\partial t} + A_0 \frac{dh_1}{dt} \eta_p^1 + A_0 \frac{d\eta_p^0}{d\tau} = A_0 \left( v_1 + x_p^1 \frac{\partial v_0}{\partial x} + \eta_p^1 \frac{\partial v_0}{\partial \eta} \right).$$

Taking the short-time average $\langle \cdot \rangle = \frac{1}{2\pi} \int_0^{2\pi} \cdot dt$, the periodic terms vanish and

$$\left\langle \frac{\partial}{\partial t} h_1 \left( \int v_0 dt - h_1 \eta_p^0 ) \right) \right\rangle = 0,$$

leaving

$$\frac{dx_p^0}{d\tau} = \langle u_1 \rangle + A_0 \left\langle \left( \int v_0 dt - h_1 \eta_p^0 \right) \frac{\partial u_0}{\partial \eta} \right\rangle,$$

$$\frac{d\eta_p^0}{d\tau} = \langle v_1 \rangle + A_0 \left\langle \left( \int v_0 dt - h_1 \eta_p^0 \right) \frac{\partial v_0}{\partial \eta} \right\rangle.$$

The Lagrangian velocity is defined as

$$u_L = \frac{dx_p^0}{d\tau} = \langle u_1 \rangle + u_\omega,$$

$$v_L = \frac{d\eta_p^0}{d\tau} = \langle v_1 \rangle + v_\omega,$$

where $\langle \boldsymbol{u}_1 \rangle = \boldsymbol{u}_s + \boldsymbol{u}_{pd}$ is the summation of the steady flows derived in the previous section, and $\boldsymbol{u}_\omega$ is the Stokes drift of the fluid particle, which is the drift resulting from the oscillatory flow.

#### S6 Parametric dependence of fluid velocity components

From the previous section, the Lagrangian velocity is comprised of a sum of the steady streaming, Stokes drift and production–drainage flow. To understand the effect of key parameters on the fluid flows, we analyse the effect of varying the amplitude of oscillations $A$, and the frequency of oscillations through the Womersley number $\alpha$.

The dimensionless steady streaming velocities depend on $A$ linearly (as it appears explicitly in the expression), verified in figure S3, and $\alpha$ quadratically for approximately $\alpha < 4$, as shown in figure figure S4. We verify that steady streaming is invariant with the scaling by plotting $\boldsymbol{u}_s/(A\alpha^2/\varepsilon)$ in figure S5. We choose appropriate ranges of $A$ and $\alpha$ based on our assumptions that $\alpha^2 \sim O(1)$ and $A \sim O(\varepsilon)$. We can see from figure S5 that independent of parameters $A$, $\alpha$, and $\varepsilon$, the steady streaming components $u_s$ and $v_s$ are scaled by constants $\gamma_u = \max(u_s)/(A\alpha^2/\epsilon)$ and $\gamma_v = \max(v_s)/(A\alpha^2/\epsilon)$, respectively. We therefore approximate the steady streaming components as $u_s \sim \gamma_u A\alpha^2/\varepsilon$ and $v_s \sim \gamma_v A\alpha^2/\varepsilon$. We

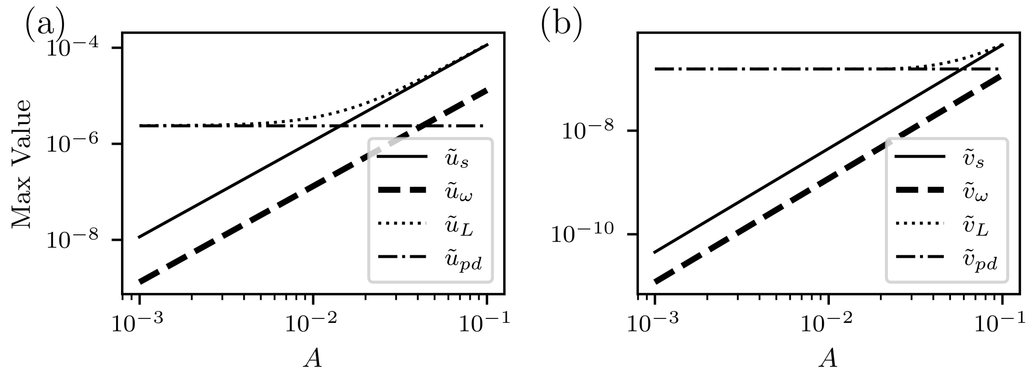


Figure S3: Comparison of dimensional velocity components (in m/s) as $A$ is varied for $\alpha = 2.5$. Figure (a) shows the horizontal velocity components and (b) the vertical velocity components.

account for this scale when defining the effective Péclet number in equation (3.1) in section 3.2 in the main text by including the coefficient $\gamma_u = \epsilon \max(u_s)/A\alpha^2$. For $\alpha < 4$, $\gamma_u = 0.006$. For larger $\alpha$ $\gamma_u$ is smaller, e.g. for $\alpha = 4.5$ used in simulation, $\gamma_u = 0.0053$. We incorporate this $\gamma_u$ in order to calculate the exact $Pe_s$.

The production–drainage flows are constant in dimensional form, and therefore solely depend on the variables introduced by the dimensionless scaling. The maximum velocity components for Lagrangian, steady streaming, Stokes drift and production–drainage as $\alpha$ and $A$ are varied are shown in figures S3 and S4 respectively. The Stokes drift has the same parametric variability as steady streaming. The magnitude of Stokes drift is always an order of magnitude less than the steady streaming, therefore Stokes drift makes a negligible contribution to the Lagrangian velocity. The steady-streaming and Stokes-drift vertical velocity components are $O(\varepsilon)$ smaller than the horizontal velocity components, as we would expect.

#### S7 Average transport equation

The transport of solutes is governed by the advection–diffusion equation (S19). Leveraging the existence of the short timescale $\tilde{t} \sim \omega^{-1}$ and the long timescale $\tilde{\tau} \sim (\varepsilon^2 A_0 \omega)^{-1}$, we perform a multiple-timescale expansion of the advection–diffusion equation to derive the time-averaged evolution of the solute. Expanding equation (S19) in multiple timescales, and letting $\sigma = \varepsilon^2 S$,

$$\frac{\partial c}{\partial t} + \varepsilon^2 A_0 \frac{\partial c}{\partial \tau} + \varepsilon A_0 u \frac{\partial c}{\partial x} + \frac{1}{h(t)} \left( \varepsilon A_0 v - h'(t)\eta \right) \frac{\partial c}{\partial \eta} = \frac{\varepsilon^2}{\alpha^2 \sigma h^2(t)} \left( \frac{\partial^2 c}{\partial \eta^2} + \varepsilon^2 \frac{\partial^2 c}{\partial x^2} \right).$$

The concentration $c$ can be expanded asymptotically as

$$c = c_0 + \varepsilon c_1 + \varepsilon^2 c_2 + O(\varepsilon^3),$$

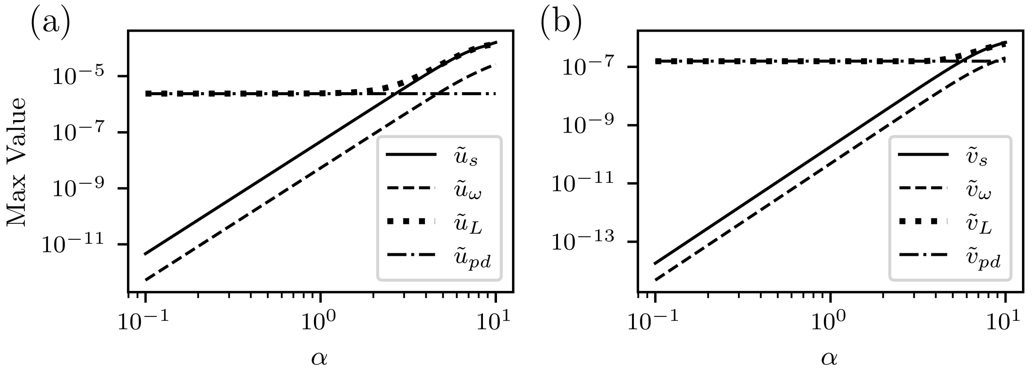


Figure S4: Comparison of dimensional velocity components (in m/s) as $\alpha$ is varied for $A = 0.0125$. Figure (a) shows the horizontal velocity components and (b) the vertical velocity components.

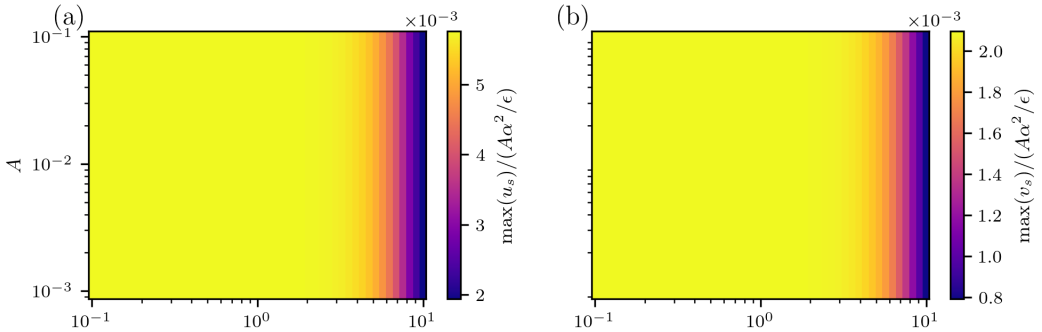


Figure S5: Maximum steady streaming velocities divided by $A\alpha^2/\varepsilon$ for varying $A$ and $\alpha$.

where each $c_i$ is comprised of both an oscillatory component $c_i^{osc}$ and a steady component $\langle c_i \rangle$. The velocities and oscillatory function $h$ are expanded using expansions (S21) and (S22) respectively. The $h^{-1}(t)$ and $h^{-2}(t)$ terms are expanded using a Taylor expansion about $\varepsilon = 0$. Then the expression up to $O(\varepsilon^2)$ is

$$\frac{\partial c}{\partial t} + \varepsilon^2 A_0 \frac{\partial c}{\partial \tau} + \varepsilon A_0 u \frac{\partial c}{\partial x} + \varepsilon A_0 \left( v - \frac{dh_1}{dt}\eta \right) \frac{\partial c}{\partial \eta} - \varepsilon^2 A_0^2 h_1 \left( v - \frac{dh_1}{dt}\eta \right) \frac{\partial c}{\partial \eta} = \frac{\varepsilon^2}{\alpha^2 \sigma} \frac{\partial^2 c}{\partial \eta^2},$$

which leads to three problems at $O(1)$, $O(\varepsilon)$, and $O(\varepsilon^2)$, respectively, which may be solved sequentially.

At leading-order:

$$\frac{\partial c_0}{\partial t} = 0 \implies c_0(x, \eta, \tau).$$

At $O(\varepsilon)$:

$$\frac{\partial c_1}{\partial t} + A_0 u_0 \frac{\partial c_0}{\partial x} + A_0 \left( v_0 - \frac{dh_1}{dt}\eta \right) \frac{\partial c_0}{\partial \eta} = 0,$$

which can be integrated directly with respect to $t$

$$c_1 = -c_{0x} A_0 \int \left( u_0 - c_{0\eta} \left( v_0 - \frac{dh_1}{dt} \right) \right) dt + \langle c_1 \rangle.$$

At $O(\varepsilon^2)$:

$$\frac{\partial c_2}{\partial t} + A_0 \frac{\partial c_0}{\partial \tau} + A_0 \Bigg( u_1 \frac{\partial c_0}{\partial x} + u_0 \frac{\partial c_1}{\partial x} + \left( v_1 - A_0 h_1 \left( v_0 - \frac{dh_1}{dt}\eta \right) \right) \frac{\partial c_0}{\partial \eta} + \left( v_0 - \frac{dh_1}{dt}\eta \right) \frac{\partial c_1}{\partial \eta} \Bigg) = \frac{1}{\alpha^2 \sigma} \frac{\partial^2 c_0}{\partial \eta^2}.$$

Taking the time average $\langle \cdot \rangle = \frac{1}{2\pi} \int \cdot dt$, the long-time transport equation averaged over the period of oscillations emerges as

$$\frac{\partial c_0}{\partial \tau} + A_0 \left( u_L \frac{\partial c_0}{\partial x} + v_L \frac{\partial c_0}{\partial \eta} \right) = \frac{1}{\alpha^2 \sigma} \frac{\partial^2 c_0}{\partial \eta^2}. \tag{S48}$$

The boundary condition (S20), taking the short-time average $\langle \cdot \rangle$ is

$$-\frac{\partial c_0}{\partial \eta} = \beta^* c_0 - A_0 \alpha^2 \sigma \frac{\langle v_1 \rangle \cdot c_0}{2}, \tag{S49}$$

at leading-order concentration. Since $\langle v_1 \rangle = q(x)$ by definition we substitute that into the boundary condition.

### S8 Convergence analysis

We investigated the convergence of the finite element method with respect to both spatial mesh resolution and time step size. The simulations were performed in the half-domain $x \in [0.5, 1]$.

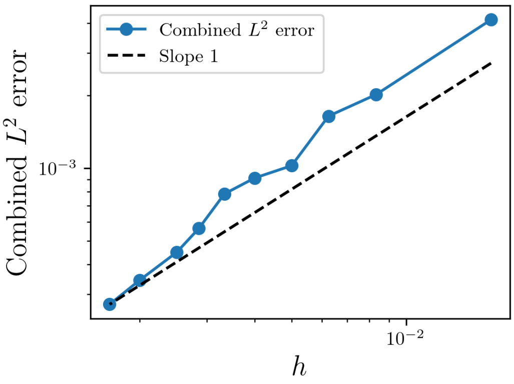


Figure S6: Spatial convergence of the finite element method for the steady-state point-leak test case with $Pe = 1$. The combined $L^2$-error was computed using concentration profiles evaluated at $\eta = -0.25$, $-0.5$, and $-0.75$, relative to a reference solution computed on a mesh with $(n_x, n_y) = (400, 800)$. The concentration and fluid fields were discretised using quadratic continuous Lagrange (CG) finite elements. The solution exhibits monotonic convergence as the mesh is refined.

To investigate spatial convergence, we considered the steady-state point-leak test case (case 2). Both the fluid and concentration fields were discretised using quadratic continuous Lagrange (CG) finite elements. Meshes of size $(n_x, n_y) = [(30, 60), (60, 120), (80, 160), (100, 200), (125, 250), (150, 300), (175, 350), (200, 400), (250, 500), (300, 600)]$ were compared against a reference solution computed on a mesh of size (400,800). To quantify the error, we compared the concentration profiles at $\eta = -0.25$, $-0.5$ and $-0.75$ with a reference solution using a discrete $L^2$-norm. For a profile sampled at points $x_i$, this was computed as

$$e_{L^2} = \left[ \sum_i \left(c(x_i) - c_{\text{ref}}(x_i)\right)^2 \Delta x \right]^{1/2},$$

where $c(x_i)$ is the concentration value at the sampled location $x_i$, and $c_{\text{ref}}(x_i)$ is the corresponding value from the reference solution. The combined error over the three $\eta$−coordinates is given by

$$e_{L^2} = \left[ \sum_{j=1}^{3} \sum_i \left(c_j(x_i) - c_{\text{ref},j}(x_i)\right)^2 \Delta x \right]^{1/2}.$$

The corresponding combined ($L^2$)-errors are shown in figure S6. The solution exhibits monotonic convergence as the mesh is refined.

To determine an appropriate balance between mesh resolution and polynomial order, we compared element orders (1)–(3) on meshes of size (100,200) and (200,400). Taking the (200,400) cubic-element solution as the reference, we computed the relative ($L^2$)-errors from

$$e_{\text{rel}} = \frac{e_{L^2}}{\|c_{\text{ref}}\|_{L^2}},$$

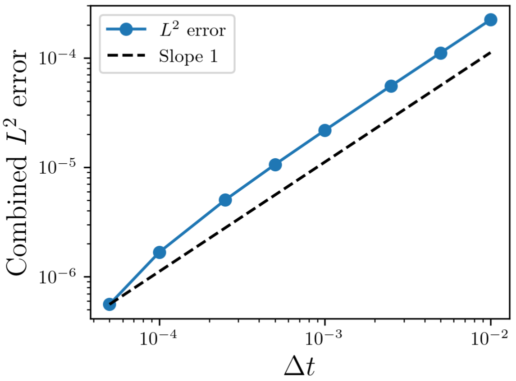


Figure S7: Temporal convergence of the finite element method for the time-dependent Gaussian initial condition test case with $Pe = 1$, using the combined $L^2$-error combined over the concentration profiles at $\eta = -0.25$, $-0.5$, and $-0.75$. Errors were computed relative to a reference solution obtained using a fine mesh with $(n_x, n_y) = (100, 200)$ and time step $\Delta t = 2.5 \times 10^{-5}$. The scheme exhibits approximately first-order convergence in time.

which are summarised in table S2. The (100,200) mesh with quadratic elements produced a relative error of only (0.26%), while remaining computationally cheaper than the finer mesh configurations. We therefore used a (100,200) mesh with quadratic elements for all production simulations.

| Mesh | Element | $L^2$ error | Relative error (%) |
|---|---|---|---|
| $100 \times 200$ | CG1 | $2.34 \times 10^{-3}$ | 0.977 |
| $100 \times 200$ | CG2 | $6.17 \times 10^{-4}$ | 0.258 |
| $100 \times 200$ | CG3 | $3.43 \times 10^{-4}$ | 0.143 |
| $200 \times 400$ | CG1 | $1.08 \times 10^{-3}$ | 0.450 |
| $200 \times 400$ | CG2 | $2.91 \times 10^{-4}$ | 0.122 |

Table S2: Comparison of mesh resolution and polynomial element order using the relative $L^2$ profile error. The reference solution corresponds to the $200 \times 400$ mesh with cubic (CG3) elements.

The implemented method uses a backward Euler time-stepping scheme. We analysed the combined $L^2$-error of the concentration profiles at $\eta = -0.25$, $-0.5$, and $-0.75$ for the time-dependent Gaussian initial condition test case with $Pe = 1$. The reference solution was generated using the mesh $(n_x, n_y) = (100, 200)$ and a very small time step $\Delta t = 2.5 \times 10^{-5}$. The combined $L^2$-errors are depicted in figure S7. The scheme exhibits approximately first-order convergence in time, consistent with the backward Euler discretisation.

Finally, we verified that the chosen time step ($\Delta t = 10^{-3}$) introduced negligible additional temporal error. Comparing the solution at ($\Delta t = 10^{-3}$) with that at ($\Delta t = 5 \times 10^{-4}$) yielded a relative ($L^2$)-error of only (0.017%), indicating that further time step refinement produces negligible changes in the solution.

#### S9 Additional results for solute transport

##### S9(a) Effect of varying $\beta_0^*$

In the results presented in section 3, we assume that $D_{AG} = 10^{-10}$ m$^2$/s. We note that while this value is not reported in the literature, varying this parameter i.e. increasing/ decreasing the permeability of the intracranial drainage site leads to qualitatively similar results. In figure S8, the solute mass and percentage drainage through arachnoid granulations for a point leak at $x_0 = 0.75$ are shown. There is a limited effect of varying $\beta_0^*$, beyond increasing mass accumulation in the case where $\boldsymbol{u}_{pd} = 0$.

##### S9(b) Case study 2b

In this section, we discuss the steady-state profiles of solute emitted uniformly at the surface of the brain. In figure 10 we show the total mass and the fraction of clearance through arachnoid granulations for different $Pe_s$ (dash-dotted magenta lines). The steady-state concentration profiles are shown in figure S9.

In the case of steady streaming alone (figure S9a), for small $Pe_s$ there is solute accumulation in the region near the spine, where there is no clearance via arachnoid granulations. As $Pe_s$ increases, more solute from this region clears into the spine, and the mass decreases. For very large $Pe_s \sim 10$, there is a small accumulation of solute in the bottom centre (around $x = 1/2$) of the domain (figure S9), where the advection components are small.

When production–drainage flow is included (figure S9b), mass accumulation is greatly decreased for lower $Pe_s$. Solutes are cleared by the arachnoid granulations, and solute accumulation near the brain at $\eta = -1$ is due to weak solute advection there.

##### S9(c) Case study 3

In this section we include some additional results for case study 3, outlined in section 3.2.3. The evolution of a solute injected intrathecally is shown in figure S10 for an intermediate steady streaming Péclet number, $Pe_s = 1$. The solute is advected into the channel by the negative longitudinal velocity about the centreline $\eta = -0.5$, and diffuses transversally towards the upper and lower boundaries, where positive velocities advect it back into the spinal canal, thus forming a trapping region. For balanced advection and diffusion, the resulting steady state depicts the front of the concentration profile reaching $x \sim 0.75$. The steady state reached varies for different $Pe_s$, as seen in figure 11, in particular the location reached by the front increases as $Pe_s$ increases.

The saturation location of the solute front, defined as the point on the surface of the brain closest to the midpoint $x = 0.5$ where $c_0 > 0.01$, is plotted in figure S11a. For $\boldsymbol{u}_{pd} = 0$,

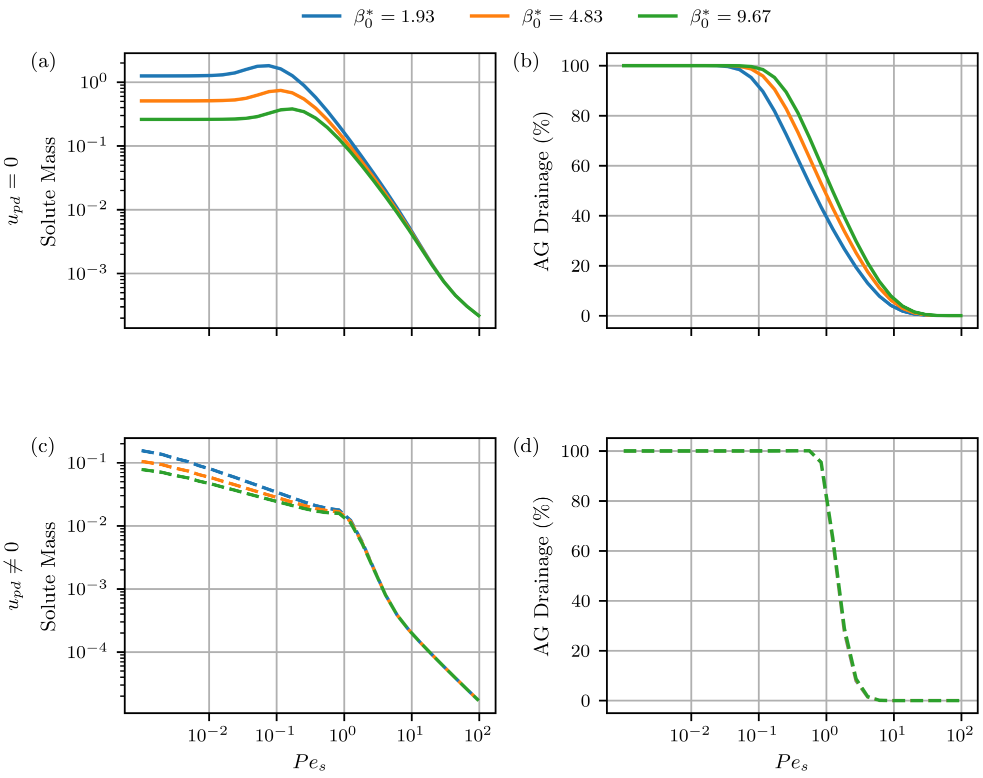


Figure S8: Plot of solute mass and percentage drainage at steady state through the arachnoid granulations for a point leak at $x_0 = 0.75$ for different $D_{AG}$. $D_{AG}$ is changed by varying the dimensionless parameter $\beta_0^* = \frac{D_{AG}}{H_{AG}} \cdot \frac{H}{D}$. For less permeable arachnoid granulations $\beta_0^* = 1.93$ corresponding with $D_{AG} = 2 \times 10^{-11}$ m$^2$/s, more solute mass accumulates in the channel, and the peak in solute mass occurs for a lower $Pe_s$ in the case of steady streaming only. $\beta_0^* = 1.93$ (blue), $\beta_0^* = 4.83$ (orange) corresponds with $D_{AG} = 5 \times 10^{-11}$ m$^2$/s used in simulations and $\beta_0^* = 9.67$ corresponds to $D_{AG} = D$. The percentage drainage through arachnoid granulations is very similar in each case, with the transition to predominant drainage through the spine occurring earlier in the case of less permeable membrane. When production-drainage flow is included, there is very little change caused by varying $D_{AG}$. The steady-state results will be qualitatively similar for different membrane permeabilities.

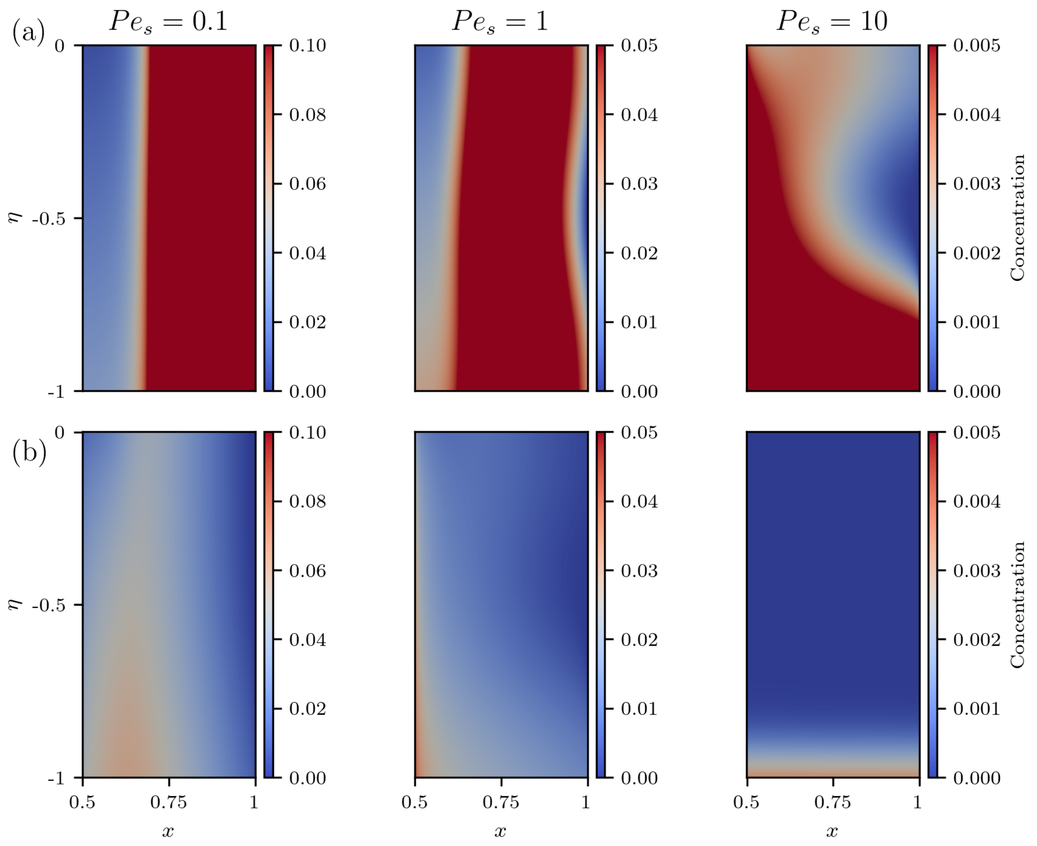


Figure S9: Steady-state concentration profiles for the full brain leak (case 2b) at different $Pe_s$ (a) with steady streaming only ($\boldsymbol{u}_{pd} = 0$) and (b) with production–drainage flows.

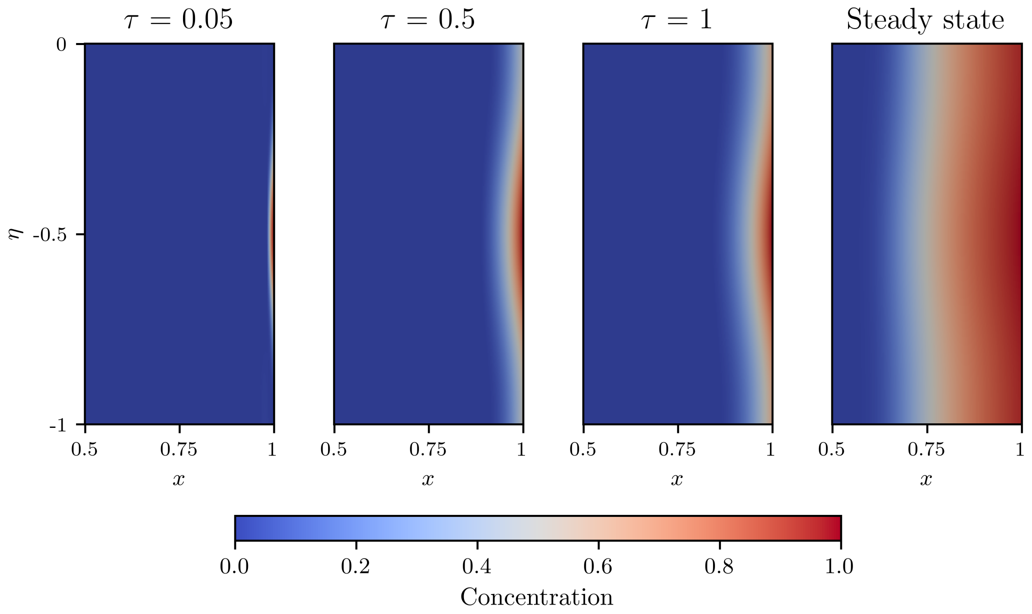


Figure S10: Example of case 3 for $Pe_s = 1$ with $\boldsymbol{u}_{pd} = 0$. The first three panels show the evolution of the solute in time. The final panel depicts the steady state reached.

a large $Pe_s$ is required for the solute to reach and saturate most of the brain surface. In contrast, when production–drainage flow is included, the solute saturates the entire surface for $Pe_s > 0.1$.

Figure S11b shows the corresponding percentage of solute cleared through arachnoid granulations. In the case of steady streaming alone, clearance occurs via the spine for low $Pe_s$, as seen in figure 11a. As $Pe_s$ increases some clearance through arachnoid granulations occurs. For $Pe_s \sim \mathcal{O}\left(10^1\right)$, clearance via arachnoid granulations reaches its peak. Beyond this, the strong outward advection counteracts the inward advection of solutes towards the granulations, reducing their contribution to clearance. When production–drainage flow is included, clearance through the arachnoid granulations is enhanced at low $Pe_s$. As the strength of steady streaming increases and becomes comparable with the production–drainage flow, the dominant clearance pathway gradually shifts toward the spinal canal. However, the inward transport enhancement due to production–drainage flow delays this transition, sustaining a higher level of granulation-mediated clearance as far as $Pe_s \sim \mathcal{O}\left(1\right)$ in comparison to the case with no production–drainage.

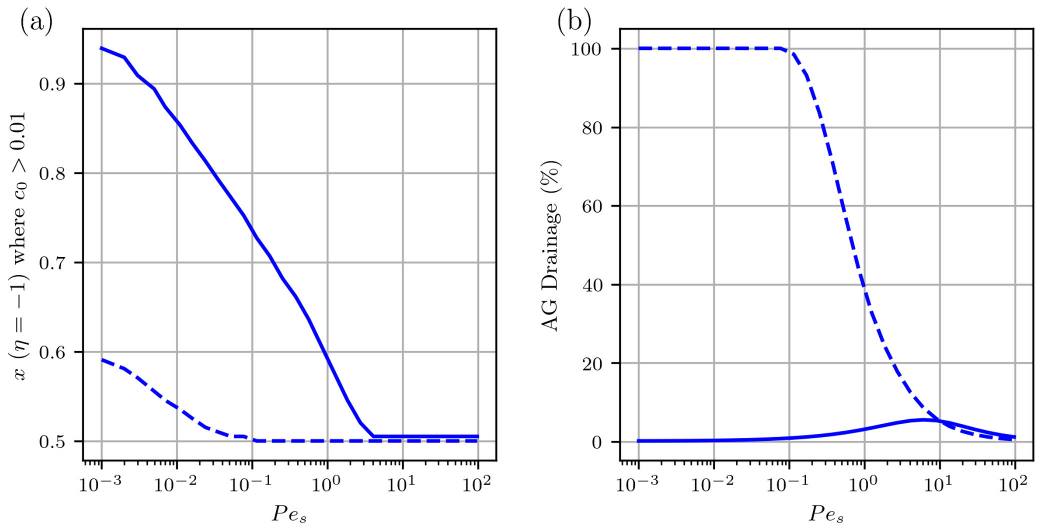


Figure S11: (a) The location at the brain surface of the concentration front exceeding 1%. (b) The percentage of drainage occurring via arachnoid granulations for a solute entering the cSAS from the spinal canal. In both panels, solid lines indicate steady streaming only and dashed lines include production–drainage flow.

### S10 Parameter Estimation

#### S10(a) Geometry

##### S10(a).1 Brain radius $R$

**Humans:** The volume of the brain can be calculated from [3] where the volumetric strain $\varepsilon_k$ is equated with the ratio between brain tissue volume change $\Delta V$ to original brain tissue volume $V$. The mean peak volumetric strain on the first scan is $\varepsilon_k = (6.4 \pm 1.7) \times 10^{-4}$ and the corresponding mean peak volume changes are $\Delta V = 0.82 \pm 0.24$ ml. Considering the mean values of $\varepsilon_k$ and $\Delta V$ the mean brain tissue volume is

$$V = \frac{\Delta V}{\varepsilon_k},$$
$$V = 1281 \text{ ml.}$$

Approximating the brain as a sphere such that $V = \frac{4}{3}\pi R^3$ we obtain the brain radius $R = 6.7$ cm.
**Mice:** The volume of a mouse brain is 415 mm$^3$ according to [4]. Assuming a spherical brain, $V = \frac{4}{3}\pi R^3$ and $R = 4.63$ mm.

##### S10(a).2 Channel length $L$

The length of the channel $L$ is defined as the circumference of the brain, such that $L = 2\pi R$.
**Humans:** Using $R$ from above, the length of the channel is $L = 2\pi R = 42.1$ cm.

**Mice:** Similarly, $L = 2\pi R = 29.1$ mm.

##### S10(a).3 Thickness of SAS $H$

**Humans:** The thickness of the cranial subarachnoid space was measured in [5] at the right frontal lobe, left frontal lobe, right occipital lobe and left occipital lobe. The difference in the case of the two hemispheres in the frontal and occipital lobes was deemed statistically insignificant. We therefore only consider the measurements from the right hemisphere, $H_{frontal} = 2.40 \pm 1.17$ mm and $H_{occipital} = 0.5 \pm 0.35$ mm. These measurements can be considered independent, and therefore to have independent standard deviation and mean. The mean height of SAS can be calculated directly from the means of the two regions as $\bar{H} = 1.45$ mm. Applying the standard law for additive error propagation we can obtain the relative uncertainty

$$(\sigma_H)^2 = \left(\frac{\sigma_F}{2}\right)^2 + \left(\frac{\sigma_O}{2}\right)^2,$$
$$\sigma_H \approx 0.61.$$

The height of the SAS is therefore $H = 1.45 \pm 0.61$ mm.
**Mice:** We estimate the height of the SAS to be $30 - 50$ $\mu$m in mice, from Figure 1 in [6].

#### S10(b) Fluid properties

CSF is a water-like Newtonian fluid [7]. CSF viscosity is $\mu = 0.8 \times 10^{-3}$ Pa·s [8] and CSF density is considered equivalent to that of water $\rho = 1007$ kg/m$^3$ [9]. The kinematic viscosity, $\nu = \mu/\rho$, is calculated from the above values to be $\nu = 0.8 \times 10^{-6}$ m$^2$/s.

#### S10(c) Production–drainage flow

##### S10(c).1 $Q_{prod}$

**Humans:** CSF production rate was measured to be between 300-1200 $\mu$l/min [10], with the commonly reported figure of 500 ml/day $\approx 5.8 \cdot 10^{-9}$ m$^3$/s being used in this study.
**Mice:** The following measurements are available: $1.5 \cdot 10^{-12}$ m$^3$/s [11], and $5.8 \cdot 10^{-12}$ m$^3$/s [12]. Here we use the more recent value from [11].

##### S10(c).2 $u_{prod}$

The velocity of CSF into the SAS is approximated by taking the volume of CSF produced, $Q_{prod}$ and dividing by the area of the SAS between the spherical brain and dura membrane shell. Taking the cross-sectional area between the spherical brain and dura membrane shell (at the equator), then $A_c = \pi\,(R+h)^2 - \pi R^2$, which gives at leading-order $A_c \approx 2\pi R h$. Then

$$\tilde{u}_{prod} = \frac{Q_{prod}}{2\pi R h},$$

The dimensionless value of $u_{prod}$ is found by multiplying the value by $\varepsilon/A\omega H$. We assume this value is the maximum velocity at the inlet.
**Humans:** $\tilde{u}_{prod} \approx 9.5 \times 10^{-6}$ m/s.
**Mice:** $\tilde{u}_{prod} \approx 9.95 \times 10^{-7}$ m/s.

### S10(c).3 $q(x)$

The (dimensionless) boundary condition $v(x, 0) = \varepsilon q(x)$ is given by

$$q(x) = q_0\left(1 - \frac{1}{1 + \exp\big(-\sigma_d(x - x_d)\big)} - \frac{1}{1 + \exp\big(-\sigma_d(1 - x - x_d)\big)}\right), \tag{S50}$$

which we abbreviate to $q(x) = q_0 d(x)$. We define the function $q(x)$ such that there is flow through the arachnoid granulae close to $x = 1/2$, and no flow through the dura membrane away from the arachnoid granulae. The velocity of CSF from production into the SAS can be approximated as parabolic flow $\tilde{u}_{in} = 4\tilde{u}_{prod}\tilde{y}(\tilde{y} + H)/H^2$, or in dimensionless form $u_{in} = 4\tilde{u}_{prod}(\varepsilon/A\omega H)y(y + 1)$. The flux into the SAS must balance the flux out through the arachnoid granulations and therefore

$$\int_{1/2}^{1} \varepsilon q(x)dx = \frac{\varepsilon}{A\omega H}\int_{-1}^{0} \tilde{u}_{in}dy,$$

which, can be arranged for an expression of $q_0$ dimensionless

$$q_0 = \frac{4\tilde{u}_{prod}}{6A\omega H \int_{1/2}^{1} d(x)dx}.$$

For sufficiently large $\sigma_d$ e.g. $\sigma_d \geq 25$ the integral can be approximated as $x_d - 1/2$.

##### S10(c).4 $A_{drain}$

We denote the area of the dura membrane through which drainage into the saggital sinus occurs as $A_{drain}$.
**Humans:** We estimate $A_{drain}$ using the distribution of arachnoid granulations in Figure 5 [13]. Taking the average length of the brain to be 167 mm [14] as a reference scale, the major $L_{AG}$ and minor $W_{AG}$ axes of the area populated by arachnoid granulations can be measured. The regions in each brain hemisphere where arachnoid granulations are likely to occur are then approximated by ellipses with major axis $L_{AG} = 15.8$ cm and minor axis $W_{AG} = 5$ cm. The area of the ellipse is calculated in metres as

$$\pi(L_{AG}/2)(W_{AG}/2) = \pi(0.025)(0.079) \tag{S51}$$
$$= 6.2 \times 10^{-3}\ \text{m}^2. \tag{S52}$$

This area would be even larger if the curvature is accounted for.

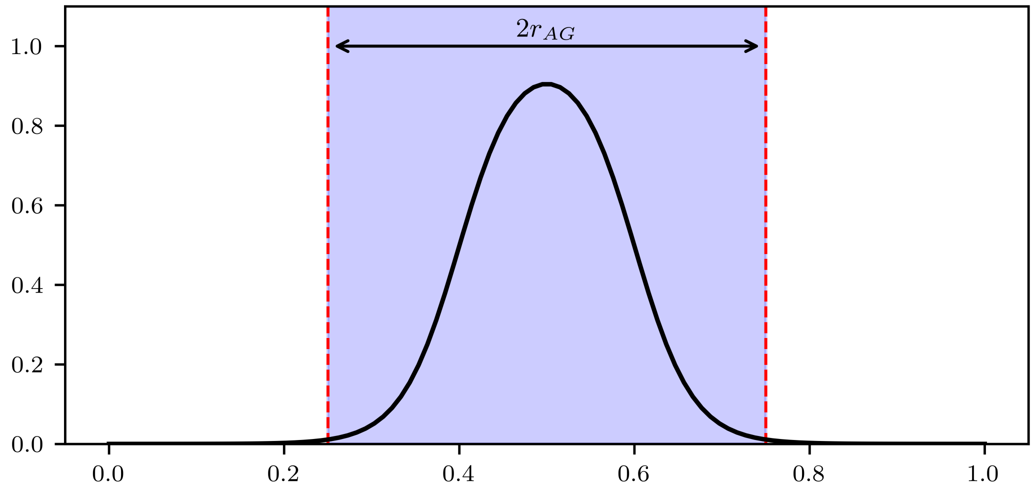


Figure S12: Plot of $q(x)$ used in simulations with $x_d = 0.6$ and $\sigma_d = 30$. The region where drainage occurs is highlighted blue, and lies within the region $x \in [0.5 - r_{AG}, 0.5 + r_{AG}]$, the boundaries of which are depicted by the red vertical lines.

###### S10(c).5 $x_d$ and $\sigma_d$

**Humans:** We assume that the area of the arachnoid granulations corresponds to a cylindrical section of the dura such that the radius of the arachnoid granulations $\tilde{r}_{AG}$ is

$$\pi \tilde{r}_{AG}^2 = A_{drain}, \tag{S53}$$

$$\tilde{r}_{AG} = \sqrt{\frac{A_{drain}}{\pi}} \tag{S54}$$

$$\approx 4.44 \times 10^{-2} \text{ m}. \tag{S55}$$

Dimensionless $r_{AG}$ is found by scaling with $L$, such that $r_{AG} = 0.105$. However, if we account implicitly for the curvature and assume that our domain spans from the top of the head to the equator, we find $\tilde{r}_{AG}/\pi(L/2) \approx 0.067$. We assume $r_{AG} = 0.1$, and that the drainage occurs approximately within the interval $1/2 - r_{AG} < x < 1/2 + r_{AG}$. To achieve this, for the function $d(x)$ we choose $x_d = 0.6$ and $\sigma_d = 30$ (see figure S12), which results in a smoother function and preserves the validity of lubrication theory.
**Mice:** We do not simulate production–drainage explicitly in mice, so no $x_d$ is calculated.

##### S10(d) Diffusion coefficients

#### S10(d).1 $D$

We choose a diffusion coefficient $D = 1.95 \times 10^{-10}$ m$^2$/s corresponding with large solutes such as amyloid-$\beta$ ($D = 1.8 \times 10^{-10}$ m$^2$/s [15]). This choice was made such that $S = \nu/D = 4096$ for round $Pe_s$ in simulations.

###### S10(d).2 $\beta^*$

$\tilde{\beta} = \tilde{\beta}_0 d(x)$, where the function $d(x)$ is the same as in equation (S50). $\tilde{\beta}_0$ is approximated by $\tilde{\beta}_0 = D_{AG}/H_{AG}$, where here we assume $D_{AG} \approx 10^{-10}$ m$^2$/s is the diffusion of the solute through the arachnoid granulations and $H_{AG} \approx 3 \times 10^{-4}$ m is the height of the arachnoid granulations [16]. We note that $D_{AG}$ has not been reported in the literature, so we make this approximation assuming that diffusion across the AG will be slower than in the unobstructed channel. Then $\tilde{\beta}_0 = 0.3 \times 10^{-6}$ m/s, and since $\beta^* = \tilde{\beta}H/D$, $\beta_0^* = 4.8$ with $\beta^* = \beta_0^* d(x)$.

#### S10(e) Oscillations

##### S10(e).1 Amplitude of cardiac pulsations

**Humans:** The amplitude of the pulsations of the brain due to the cardiac cycle are estimated from the displacement of CSF throughout the cardiac cycle in [3]. The brain is estimated as a sphere with time dependent volume, so that the volume of the cranial SAS is

$$V(t) = \frac{4}{3}\pi \left(R^3 - (R - \tilde{h}(t))^3\right),$$

where we assume $\tilde{h}(t) = H + \tilde{A}\sin(\omega\tilde{t})$ describes the time-dependent height of the SAS, and both $\tilde{A}$ and $H$ are considered to be small parameters which are negligible at $O(\tilde{A}^2)$ and $O(H^2)$. Expanding the cubic terms and retaining the leading-order terms, we obtain

$$V(t) \approx 4\pi R^2 \tilde{h}(t). \tag{S56}$$

Considering the change in height $\Delta\tilde{h} = 2\tilde{A}$, then amplitude is

$$\tilde{A} = \frac{\Delta V}{8\pi R^2},$$
$$\tilde{A} = (6.6 \pm 2.9) \times 10^{-6}\text{ m},$$

where $\Delta V = 0.74 \pm 0.33$ ml is the mean peak CSF stroke volume (measured at C2/C3), and since $\Delta V$ is the only variable carrying experimental uncertainty the standard deviation of $\tilde{A}$ is obtained by first-order error propagation.

**Mice:** We can estimate the brain volume change induced by cardiac vascular pulsations in the mouse from the relation between pressure and volume obtained by infusion tests [17]. Compliance $C$ links a change of pressure to a change of volume via $\Delta V = C\,\Delta P$. With $C = 1.798 \pm 0.185$ $\mu$L/mmHg and a typical ICP pulse amplitude at the cardiac time scale of $\Delta P \approx 1$ mmHg, the corresponding volume change is $\Delta V \approx 1.7$ $\mu$L $= 1.7 \times 10^{-9}$ m$^3$ [17]. The amplitude of cardiac oscillations in mice is calculated in the same way as in humans,

$$\tilde{A} = \frac{\Delta V}{8\pi R^2},$$
$$\tilde{A} = (3.06 \pm 0.333) \times 10^{-6}\text{ m},$$

for a brain radius of 4.7 mm. Then $\tilde{A} = 3.06 \pm 0.333$ $\mu$m.

In [18] they estimate tissue velocity using functional ultrasound (fUS). Taking 0.2-0.3 mm/s as maximum tissue velocity (from Fig 1 in that paper), we can calculate the maximum tissue displacement as $v_{max}/\omega$ = 3-5 $\mu$m using $\omega = 10 \cdot 2\pi$ Hz. This displacement is in agreement with our estimate from ICP.

##### S10(e).2 Amplitude of respiratory pulsations

**Humans:** The respiratory cycle modulates brain pulsations [19–22]. In [22], real time phase contrast MRI is used to quantify the relative contributions of respiratory and cardiac pulsations. We are interested in the flows generated by cardiac and respiratory pulsations individually, therefore we consider the measurements in this paper. Taking the ratio of the max human CSF velocity due to respiratory-only and cardiac-only pulsations (Table 1 [22]), which is $0.51/0.97 = 0.526$ for normal breathing, we scale our cardiac amplitudes by 0.526 to obtain the amplitudes of respiration induced pulsations $\tilde{A} = 3.47 \pm 1.525$ $\mu$m, which is the parameter range we consider in this work.

The authors of [21] report that the volumetric strain of brain tissue (which we know to be proportional to the amplitude of brain pulsations from our calculations in the previous section) associated with cardiac pulsations are approximately three times that of respiratory pulsations. This implies that the amplitude of respiratory pulsations would be approximately a third of those of cardiac oscillations. Considering the range of amplitudes of cardiac pulsations we calculated, the corresponding range of respiratory oscillations would be $(1.233 - 3.167)$ $\mu$m which is in good agreement with the above values.

We note that in [19, 20] the amplitudes of respiratory pulsations are not separated from the amplitudes of cardiac pulsations. To verify that our respiratory amplitude estimate is consistent, we calculate the amplitudes from these papers where cardiac and respiratory components are combined. The amplitude of these oscillations is estimated from CSF stroke volumes measured at the C2/C3 vertebral level during the respiratory cycle, as reported by [20]. The stroke volumes $V_s$ range from $0.54 - 1.66$ (ml). The stroke volume at C2/C3 is defined in [20] as

$$V_s = 0.5 \int_0^T |Q(t)| dt, \tag{S57}$$

where $T$ is the timescale of the respiratory cycle.

Recall that the leading-order approximation of time-dependent brain volume is given by (S56). The total CSF flux is given by

$$Q = \frac{dV}{dt} \sim 4\pi R^2 \frac{d\tilde{h}(t)}{d\tilde{t}},$$

and substituting $\tilde{h}(t) = H + \tilde{A}\sin(\omega\tilde{t})$ CSF stroke volume is

$$V_s = 0.5 \left(4\pi R^2\right) \int_0^T | - \tilde{A}\omega\cos(\omega\tilde{t})| d\tilde{t}. \tag{S58}$$

Since $\cos(\omega\tilde{t})$ is symmetric and periodic over $[0, T]$, the integral of its absolute value over one period can be split into four identical intervals of length $T/4$. Thus

$$\int_0^T |-\tilde{A}\omega\cos(\omega\tilde{t})|d\tilde{t} = 4\int_0^{T/4} \tilde{A}\omega\cos(\omega\tilde{t})d\tilde{t}.$$

Substituting this into (S58) and since the frequency of the respiratory cycle is $\omega = 2\pi/T$, we obtain

$$V_s = \left(4\pi R^2\right) \cdot 2\tilde{A} \sim 4\pi R^2 \Delta\tilde{h}.$$

Rearranging to isolate $\tilde{A}$, and substituting the mean stroke volume $V_s = 0.99$ ml or $0.99 \times 10^{-6}$ m$^3$, the amplitude of respiratory oscillations is $\tilde{A} = 8.775$ $\mu$m, with a range of $[4.786 - 14.71]$ $\mu$m.

In [19] the authors report flow magnitudes (ml/s) averaged over the time frames for CSF at C2/C3 for both normal and forced breathing patterns. Assuming that this quantity is $\langle|Q|\rangle = \frac{1}{T}\int_0^T |Q|dt = \frac{4\pi R^2}{T}\int_0^T |\tilde{A}\omega\cos(\omega\tilde{t})|d\tilde{t}$, we can derive the expression for $A = \langle|Q|\rangle/(8R^2\omega)$. The frequency of respiratory oscillations is taken to be in the range $f = 0.2 - 0.3$ Hz, where $\omega = 2\pi f$. The reported average flow magnitude range of $0.83 \pm 0.36$ ml/s for normal breathing rates is therefore equivalent to a respiratory oscillation of amplitude $\tilde{A} \in [3.315 - 8.394]$ $\mu$m.

We then compare the values calculated from [19, 20], where the cardiac pulsations are not separated from respiratory pulsations, with the total velocity of CSF (with the components unseparated) over the respiratory cycle from Table 1 in [22]. The amplitude of oscillations attributed to normal breathing would be $\tilde{A} = 9.385 \pm 4.12$ $\mu$m in this case, which is similar to the amplitudes calculated from [19, 20].

**Mice**: We found no measurements of CSF or brain displacement induced by respiratory pulsations in mice.

##### S10(e).3 Amplitude of sleep pulsations

**Humans:** In [23], the intensity of brain pulsations was measured using MREG and EEG during wake and sleep, and filtered by frequency of pulsations. The range of intensity of slow wave pulsations during sleep was $\approx 1500$ a.u. (Figure 2 e), while that of cardiac pulsations was $\approx 200$ a.u. Assuming direct proportionality between intensity and magnitude of oscillations, the amplitude of sleep pulsations is 7.5 times that of cardiac oscillations $\sim 49.5 \pm 21.75$ $\mu$m.

**Mice:** The volume of blood in the mouse brain ($V_{CBV}$) is measured as $5.8 \pm 0.4\%$ of total brain volume in [24] (Table 1). The brain volume of a mouse is 415 mm$^3$ ($4.15 \times 10^{-7} m^3$) [4]. Then $V_{CBV} = (2.4 \pm 0.166) \times 10^{-8}$ m$^3$.

The change in diameter of the penetrating arteriole lumen during NREM sleep was measured in [25]. The median radius of the penetrating arteriole lumen was measured as approximately 5 $\mu$m (Figure 2 f), and changes in radius of $\approx 0.5$ $\mu$m were observed (Figure 2 c) for very low frequency oscillations ($0.1 - 0.3$ Hz). The percentage change in vessel wall radius is therefore 10%. Assuming the change in vessel wall is proportional to the change in total cerebral blood volume, then $\Delta V_{CBV} = 2.4 \pm 0.166 \times 10^{-9}$ m$^3$. Equating the change in cerebral blood volume $\Delta V_{CBV}$ with the change in total brain volume $\Delta V_{brain}$, we can calculate the amplitude of these oscillations using the same method,

$$\tilde{A} = \frac{\Delta V_{CBV}}{8\pi R^2},$$
$$\tilde{A} = (2.07 \pm 0.166) \times 10^{-5} \text{ m},$$

meaning $\tilde{A} = 20.7 \pm 1.66$ $\mu$m.

We correct this for the fraction of vessels which pulsate. We excluded pial arteries because we just wanted to look at brain pulsations (though change in pial arteries will likely drive the CSF flow in cSAS). At the high end is arterioles + capillaries 59.5 ± 6%.The low end is just penetrating arterioles 2.9 ± 1.3% from [26]. Then the range of $\tilde{A} = [0.3046, 14.6065]$ $\mu$m.

##### S10(e).4 Frequency of cardiac pulsations

**Humans:** The human resting heart-rate is typically in the range 60-100 bpm [27], with a corresponding frequency of 1-1.6 Hz.
**Mice:** Mice heart-rates are typically in the range of 500-700 bpm [28], which corresponds to a frequency of $8.33 - 11.67$ Hz.

##### S10(e).5 Frequency of respiratory pulsations

**Humans:** The frequency of respiration in humans is 0.2-0.3 Hz [29], or 0.19 ± 0.11 Hz in [22]. We use the second value for consistency with our amplitude estimate.
**Mice:** The respiration rate in mice was taken to be 254.8 ± 77.8 (breaths per min) taking the mean of the values in Table 1 in [30]. We therefore consider the frequency to be in the range [2.95-5.5] Hz.

##### S10(e).6 Frequency of vasomotion pulsations

**Humans:** The frequency of vasomotion pulsations during sleep are reported as 0.05-0.1 Hz [23].
**Mice:** The frequency of very low frequency pulsations of the penetrating arterioles is reported as 0.1-0.3 Hz in [25].
